# COVID-19 virtual patient cohort reveals immune mechanisms driving disease outcomes

**DOI:** 10.1101/2021.01.05.425420

**Authors:** Adrianne L. Jenner, Rosemary A. Aogo, Sofia Alfonso, Vivienne Crowe, Amanda P. Smith, Penelope A. Morel, Courtney L. Davis, Amber M. Smith, Morgan Craig

## Abstract

To understand the diversity of immune responses to SARS-CoV-2 and distinguish features that predispose individuals to severe COVID-19, we developed a mechanistic, within-host mathematical model and virtual patient cohort. Our results indicate that virtual patients with low production rates of infected cell derived IFN subsequently experienced highly inflammatory disease phenotypes, compared to those with early and robust IFN responses. In these *in silico* patients, the maximum concentration of IL-6 was also a major predictor of CD8^+^ T cell depletion. Our analyses predicted that individuals with severe COVID-19 also have accelerated monocyte-to-macrophage differentiation that was mediated by increased IL-6 and reduced type I IFN signalling. Together, these findings identify biomarkers driving the development of severe COVID-19 and support early interventions aimed at reducing inflammation.

**Author summary:** Understanding of how the immune system responds to SARS-CoV-2 infections is critical for improving diagnostic and treatment approaches. Identifying which immune mechanisms lead to divergent outcomes can be clinically difficult, and experimental models and longitudinal data are only beginning to emerge. In response, we developed a mechanistic, mathematical and computational model of the immunopathology of COVID-19 calibrated to and validated against a broad set of experimental and clinical immunological data. To study the drivers of severe COVID-19, we used our model to expand a cohort of virtual patients, each with realistic disease dynamics. Our results identify key processes that regulate the immune response to SARS-CoV-2 infection in virtual patients and suggest viable therapeutic targets, underlining the importance of a rational approach to studying novel pathogens using intra-host models.

## Introduction

Clinical manifestations of SARS-CoV-2 infection are heterogeneous, with a significant proportion of people experiencing asymptomatic or mild infections that do not require hospitalization. In severe cases, patients develop coronavirus disease (COVID-19) that may progress to acute respiratory distress syndrome (ARDS), which is frequently accompanied by a myriad of inflammatory indicators [1]. Mounting evidence points to a hyper-reactive and dysregulated inflammatory response characterized by overexpression of pro-inflammatory cytokines (cytokine storm) and severe immunopathology as specific presentations in severe COVID-19 [2–6]. An over-exuberant innate immune response with larger numbers of infiltrating neutrophils [7,8] arrests the adaptive immune response through the excessive release of reactive oxygen species that leads to extensive tissue damage and depletion of epithelial cells [9]. In addition, lymphopenia, in particular, is one of the most prominent markers of COVID-19 and has been observed in over 80% of patients [6, 10–12]. However, the immune mechanisms that lead to disparate outcomes during SARS-CoV-2 infection remain to be delineated.

Cytokines are critically important for controlling virus infections [13, 14] and are central to the pathophysiology of COVID-19, sometimes playing a detrimental role in the context of a cytokine storm [10]. For example, interleukin-6 (IL-6) can stimulate CD8^+^ T cell expansion under inflammatory conditions [15]; however, in hospitalized SARS-CoV-2 patients with lymphopenia, IL-6 has been shown to be elevated [16] without an increase in CD8^+^ T cell counts [17]. Type I interferons (such as IFNs-α, β [18]) also play a major role in limiting viral replication by inducing a refractory state in susceptible and infected cells [19–21]. Due to this, it has been suggested that a delay in mounting an effective IFN response may be responsible for COVID-19 severity [22] as it is for other highly pathogenic coronavirus (i.e. SARS-CoV and MERS) infections [13]. Overall, patients with severe COVID-19 present with lymphopenia [14, 23], and are likely to have increased inflammatory cytokines such as IL-6, granulocyte-macrophage colony-stimulating factor (GM-CSF), and granulocyte colony-stimulating factor (G-CSF) [7, 17, 24].

Because identifying which immune mechanisms lead to divergent outcomes can be difficult clinically, and experimental models and longitudinal data are only beginning to emerge, theoretical explorations are ideal [25]. Quantitative approaches combining mechanistic disease modelling and computational strategies are being increasingly leveraged to investigate inter- and intra-patient variability by, for example, developing virtual clinical trials [26–28]. More recently, viral dynamics models [29, 30] have been applied to understand SARS-CoV-2 within-host dynamics and their implications for therapy [31–36]. However, there are few comprehensive models that integrate detailed immune mechanisms and allow interrogation of the dynamics controlling divergent outcomes, and none have attempted to quantify the high degree of variability in patient responses to SARS-CoV-2 through modelling.

In this study, we developed a mechanistic mathematical model to describe the within host immune response to SARS-CoV-2. We explicitly modelled the interactions between epithelial cells, innate and adaptive immune cells and cytokines. The model was fit to various *in vitro, in vivo*, and clinical data, analyzed to predict how early infection kinetics facilitate downstream disease dynamics, and used to create a virtual patient cohort with realistic disease courses. Our results suggest that mild and severe disease are distinguished by the rates of monocyte differentiation into macrophages and of IFN production by infected cells. In our virtual cohort, we found that severe COVID-19 responses were tightly correlated with a delay in the peak IFN concentration and that a large increase in IL-6 was the dominate predicator of CD8^+^ T cell depletion in our virtual cohort. Importantly, these results provide insight into differential presentations of COVID-19 by identifying key regulators of severe disease manifestation particularly related to monocyte differentiation and IL-6 concentrations.

## Results

### Modelling the immune response to SARS-CoV-2 and the impact of delayed IFN on infection dynamics

To study the dynamics of SARS-CoV-2 infection and the development of COVID-19, we constructed a computational biology model of host-pathogen interactions (**Eqs. S1-S22**, with variables and parameters summarized in **Table S1** and schematic in **Figure 1**). The model includes susceptible lung epithelial cells (*S*) that encounter virus (*V*) and become infected (*I*) before turning into damaged or dead cells (*D*) due to viral infection or immune involvement. The immune response is orchestrated by a myriad of cytokines that act to stimulate the immune cell subsets present in the tissues and recruit cells from the bone marrow and circulation (**Figure 1A**). Upon infection, cells begin secreting type I IFNs (*F*) that cause lung epithelial cells to become resistant to infection (*R*) and decrease the production of newly infected cells [37]. Through stimulation by infected and dead cells, alveolar (lung tissue-resident) macrophages (*M*_*ΦR*_) become inflammatory macrophages, which also arise through macrophage (*M*) differentiation by stimulation by GM-CSF (*G*) or IL-6 (*L*) [38]. Neutrophils (*N*) are recruited to the infection site by G-CSF and release reactive oxygen species (ROS) causing bystander damage to infected and susceptible cells [39, 40]. CD8^+^ T cells (*T*) are subsequently recruited to the infection site following a delay to account for antigen presentation, with expansion modulated by type I IFN and IL-6 concentrations. See **Materials and Methods** for a complete description.

**Figure 1.**
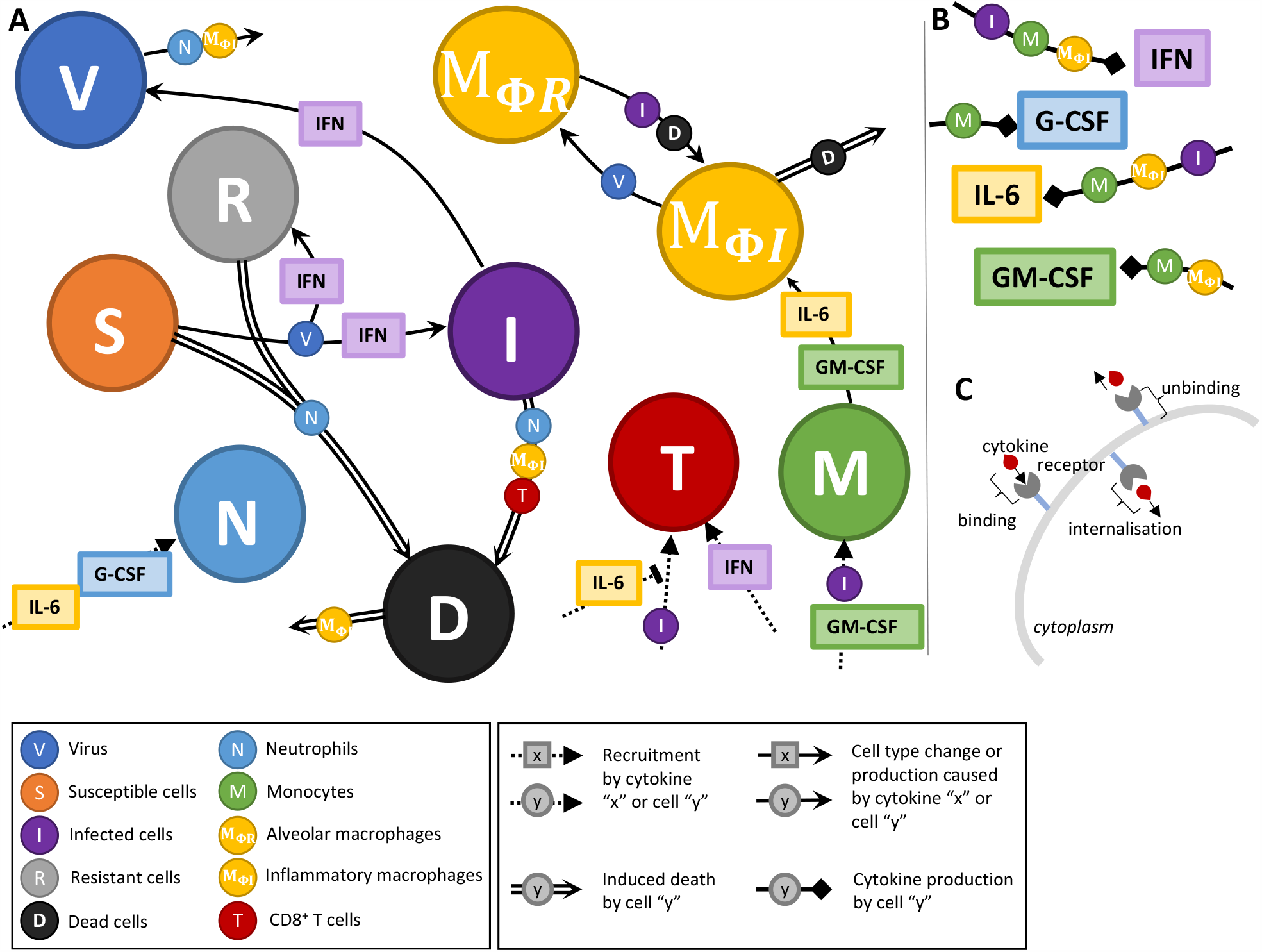
Immune response to SARS-CoV-2 infection model schematic. The model in **Eqs. S1-S22** reduced to **A)** cell dynamics **B)** cytokine production dynamics and **C)** cytokine binding kinetics. Unique lines represent induced cell death (double line), recruitment (dashed line), cell type change or production (solid line), and cytokine production (square arrow). Cell and/or cytokines along joining lines denote a causal interaction. **A)** Virus (*V*) infects susceptible lung epithelial cells and creates either infected (*I*) or resistant (*R*) cells depending on the concentration of type I IFN. Infected cells then either die and produce new virus or are removed via inflammatory macrophages (*M*_*ΦI*_) or CD8^+^ T cells (*T*) that induce apoptosis to create dead cells (*D*). Neutrophils (*N*) cause bystander damage (death) in all epithelial cells and are recruited by individually G-CSF and IL-6 concentrations. CD8^+^ T cells are recruited by infected cells and their population expands from IFN signalling. T cell recruitment is inhibited by IL-6 concentrations. Monocytes (*M*) are recruited by infected cells and GM-CSF and differentiate into inflammatory macrophages based on the individual concentrations of GM-CSF and IL-6. Tissue-resident macrophages (*M*_*ΦR*_) also become inflammatory macrophages through interaction with dead and infected cells. Dead cells are cleared up by inflammatory macrophages and also cause their death. **B)** Type I IFN is produced by infected cells, inflammatory macrophages and monocytes. G-CSF is produced solely by monocytes and GM-CSF is produced by monocytes and macrophages. IL-6 is produced by monocytes, inflammatory macrophages and infected cells. **C)** Cytokine receptor binding, internalization and unbinding kinetics considered for each cell-cytokine interaction.

Because the model has several parameters that are undetermined biologically and insufficient data exists to confidently estimate their values, we used a stepwise approach to parameter estimation (see **Materials and Methods** and **Figures S1-S5**). We first confirmed that we could recapitulate early infection viral kinetics with a reduced version of the full model (‘viral model’). For this, we excluded immunological variables (i.e. only including **Eqs. 6-9**) and estimated parameters relating to viral kinetics by fitting to viral load data from macaques (see **Materials and Methods**). The resulting model dynamics were in good agreement to these early infection data (**Figure 2**) and demonstrate a rebound in epithelial lung tissue as the viral load and infected cells decrease.

**Figure 2.**
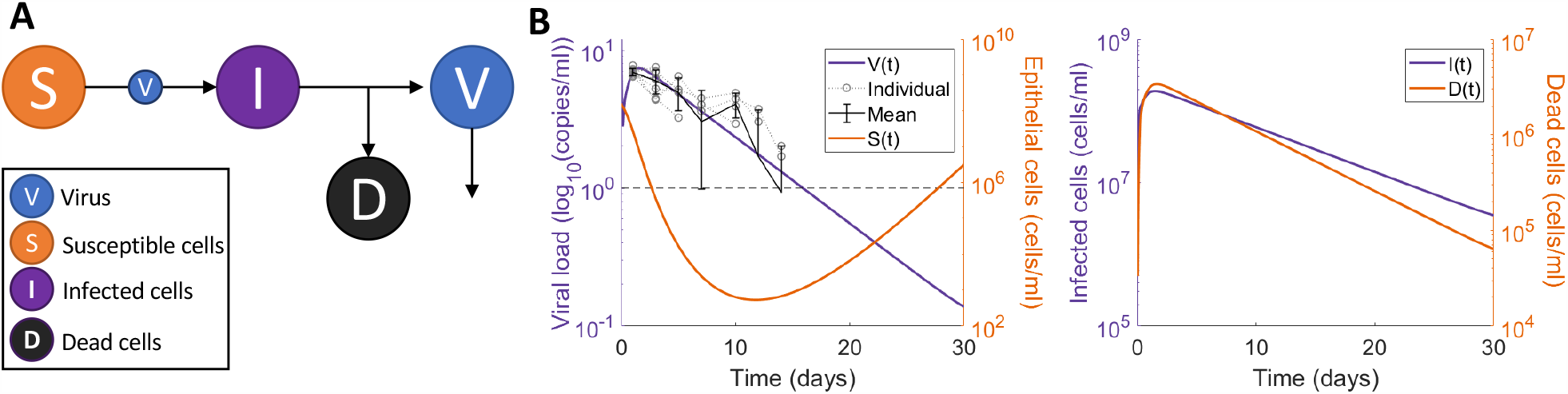
Viral dynamics model fit to macaque viral data from Munster et al. [41] A reduced version of the full model (all cytokine and immune cells set to 0, **Eqs. 6-9**) was fit to data from macaques [41] to estimate preliminary viral kinetic parameters. **A)** Virus (*V*) infects susceptible cells (*S*) making infected epithelial cells (*I*) which then die to produce dead cells (*D*) and new virus. **B)** Comparison of predicted viral dynamics compared to observations from 6 animals, with susceptible cell kinetics (left) with predictions of infected and dead cells over time (right). We estimated *β, p, d*_*I*_, *V*_0_ and *d*_*V*_ from the reduced model in **A)** fit to data from Munster et al. [41] measuring the viral load in macaques after challenge with SARS-CoV-2 (**Table S1**).

We then isolated the IFN dynamics to assess clinical and experimental findings suggesting that delaying IFN results in more severe presentations in highly pathogenic coronavirus infections including SARS-CoV-2 [13, 14, 22]. Using the parameters obtained from the ‘viral model’ (**Eqs. 6-9; Table S1**), we then simulated the impact of IFN with the ‘IFN model’ (**Eqs. 10-16** and **Figure 3A**). We examined the predicted dynamics in response to delayed IFN by simulating with and without a fixed delay for IFN production from infected cells. Our results suggest that delaying type I IFN production by 5 days yields a 10-fold increase in tissue damage with only 20% of the lung tissue remaining on day 2 (**Figure 3B**), caused by the increase in infected cells and subsequent lack of resistant cells. IFN dynamics were matched to systemic IFN-α concentrations from clinical cohorts by visual predictive check to confirm that predictions fell within the observed ranges [42] (**Figure S6**A).

**Figure 3.**
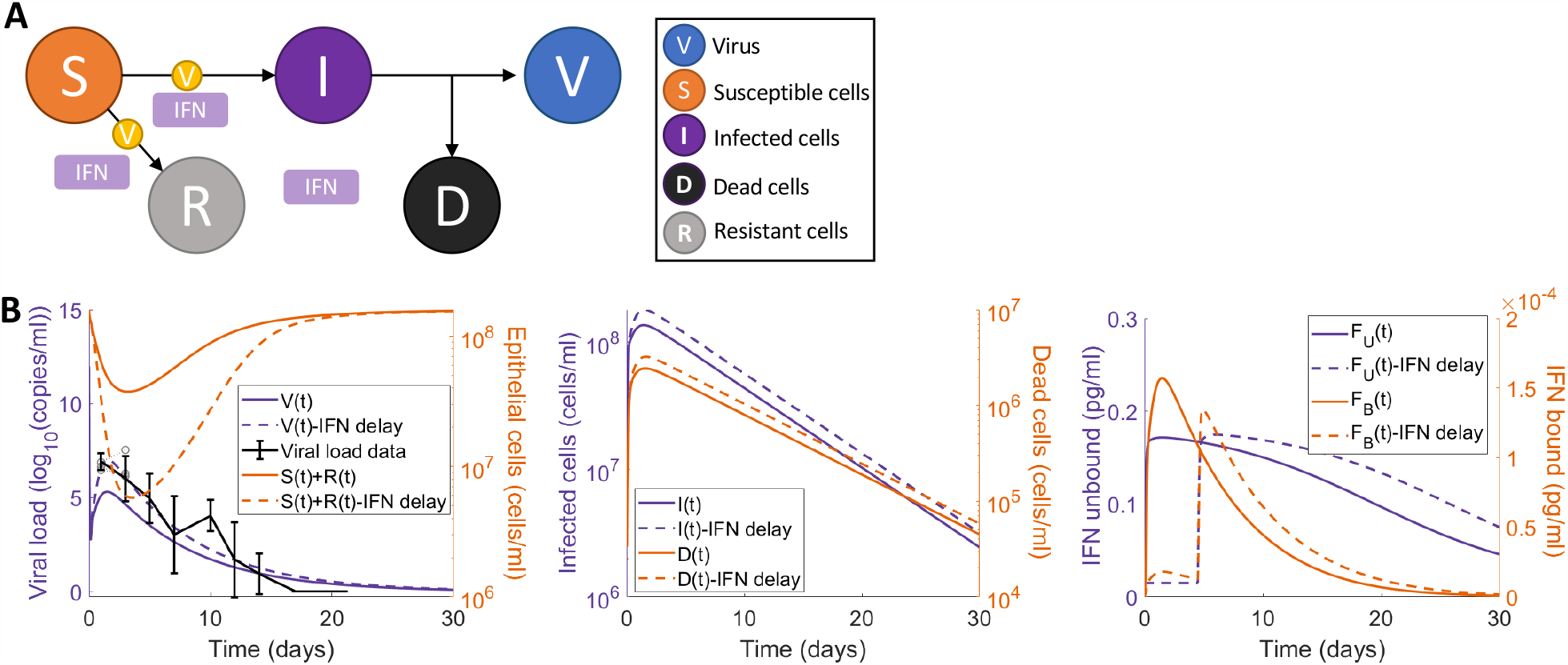
Delayed type I IFN response impacts heavily on tissue survival in reduced model. **A)** Submodel (**Eqs. 10-16**) with all non-IFN cytokines and immune cell interactions set to zero and only considering interactions between virus (*V*) and susceptible (*S*), infected (*I*), resistant (*R*), and dead (*D*) epithelial cells. **B)** Predictions from the simplified model without delayed IFN production (solid lines) versus with a constant delay (*τ*_*F*_ = 5 days) (dotted lines). Solid black (left panel): viral loads from SARS-CoV-2 infection in macaques by Munster et al. [41] is overlayed with predicted viral dynamics.

### Immunologic determinants of mild and severe disease

Next, to establish the mechanisms that differentiate mild versus severe disease, we simulated the full model (**Eqs. S1-S22**) using two different parameter sets. Mild disease dynamics were recreated using the estimated parameter values (**Table S1**) with the virus decay rate (*d*_*V*_) and the infected cell death rate (*d*_*I*_) recalculated to ensure that the maximum death rate of the virus and infected cells did not exceed the value obtained from the reduced viral dynamics model fit (**Figure 2**). Simulating mild disease, we predicted that all cell populations and cytokines rapidly return to homeostasis, with the immune response effectively clearing virus within 10 days (**Figure 4** and **Figure S7**).

**Figure 4.**
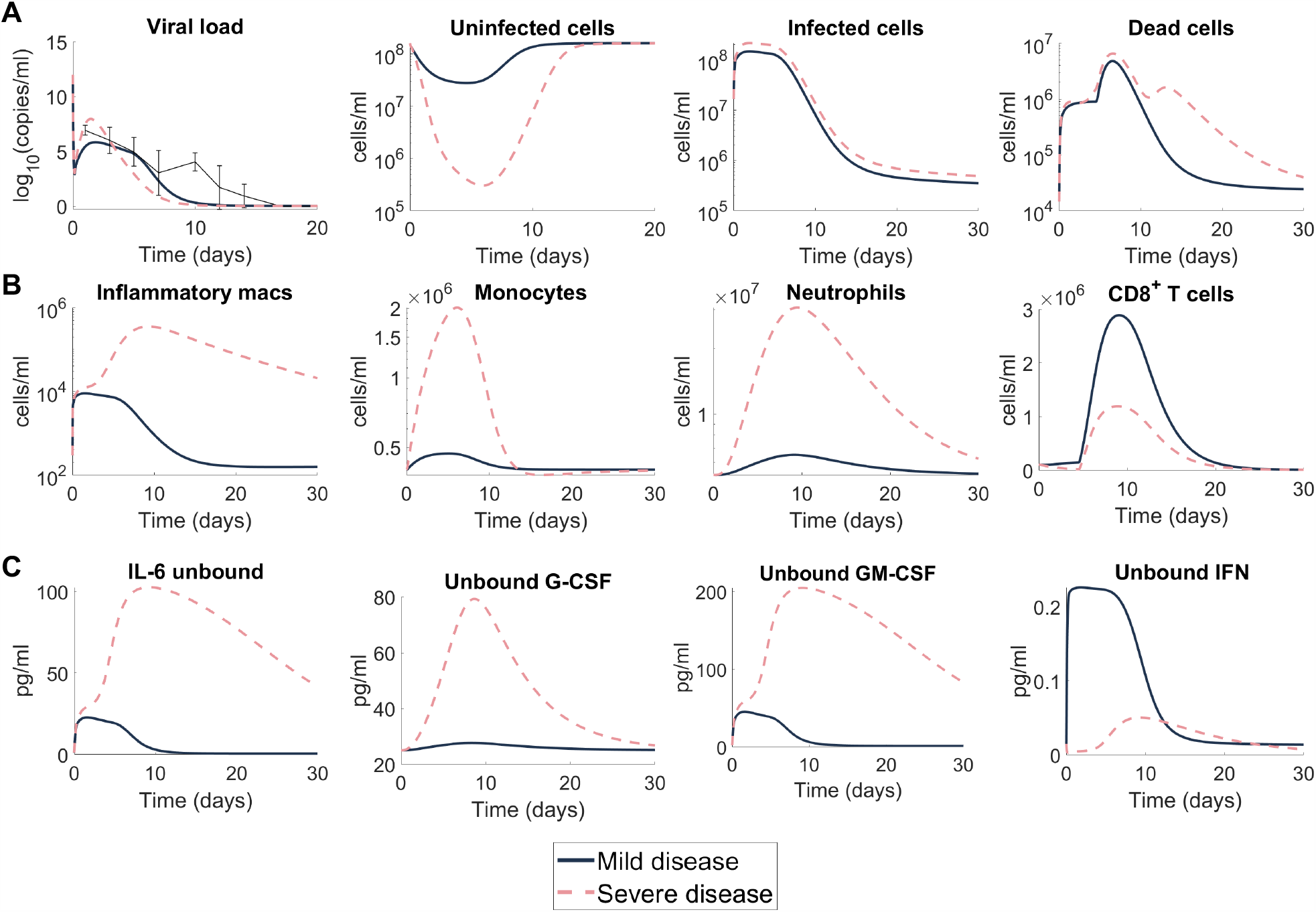
Predicting mild and severe COVID-19 dynamics. Mild disease (solid lines) dynamics obtained by using baseline parameter estimates (**Tables S1**) while severe disease dynamics (dashed lines) were obtained by decreasing the production rate of type I IFN (*p*_*F,I*_) and increasing the production of monocytes (*p_M,I_*) and their differentiation to macrophages (*η*_*F,MΦ*_). **A)** Viral load and lung cells concentrations (susceptible, resistant, infected, and dead cells). Solid black line with error bars indicates macaque data [41] (see **Figure 2**). **B)** Immune cell concentrations (inflammatory macrophages, monocytes, neutrophils, and CD8^+^ T cells. **C)** Unbound cytokine concentrations (free IL-6, GM-CSF, G-CSF, and type I IFN). Time evolution of all model variables is shown in **Figure S7** (including bound cytokine and alveolar macrophages).

Because severe SARS-Cov-2 infection results in lower levels of IFN [42] and increased monocytes [43], we recapitulated severe disease by modulating model parameters relating to these processes, i.e., the rates of IFN production from infected cells and macrophages were decreased, and the rate of monocyte recruitment from the bone marrow by infected cells was increased. With these changes, the model predicted a dramatic shift in disease response that was characterized by a cytokine storm (elevated IL-6, GM-CSF and G-CSF), high ratios of innate to adaptive immune cells, and a marked reduction in healthy viable lung tissue (**Figure 4A**), whereas changes in viral load remained relatively consistent with mild disease.

In addition, there was a significant increase in the number of inflammatory macrophages (**Figure 4B**), IL-6, GM-CSF and, importantly, a delayed and reduced IFN peak (**Figure 4C**). In comparison to the mild disease, inflammatory macrophages and neutrophils (**Figure 4B**) remained elevated for at least 30 days after initial infection. Comparing mild and severe disease highlighted significant differences in the area under the curve (AUC) of macrophages (6 × 10^4^ cells/ml versus 3 × 10^11^ cells/ml) and neutrophils (2 × 10^8^ cells/ml versus 3 × 10^13^ cells/ml) over 30 days.

Interestingly, inflammation remained high in the severe disease scenario despite the virus being cleared slightly faster (∼1 day) than in the case of mild disease (**Figure 4A**). Further, the peak of inflammatory macrophages increased from ∼10^4^ cells/ml to ∼10^6^ cells/ml in severe scenarios compared to mild scenarios (**Figure 4B**). The model also accurately predicted that CD8^+^ T cell dynamics were lower in severe cases, which is indicative of lymphopenia and similar to clinical observations from patients with severe COVID-19 [14, 23]. Despite varying only three parameters to generate disparate dynamics, the immune cell and cytokine dynamics were qualitatively in line with clinical observations for IFN-α [42], IL-6 [42, 44], and G-CSF [24] (**Figure S6**B-F).

### Macrophages, CD8^+^ T cells, IFN and IL-6 regulates response to SARS-CoV-2 infection

To further understand how the host immune system regulates the response to SARS-CoV-2 infection, we conducted a local sensitivity analysis by varying each parameter individually by ±20% and comparing a set of metrics (see **Materials and Methods**) chosen to provide a comprehensive understanding of each parameter’s impact on the host-pathogen dynamics. This analysis identified 17 sensitive parameters (**Figure 5**) relating to virus productivity (*p, δ*_*V,N*_, *β, ϵ*_*F,I*_), CD8^+^ T cell induced epithelial cell apoptosis (*δ*_*I,T*_), macrophages, monocyte and CD8^+^ T cell production 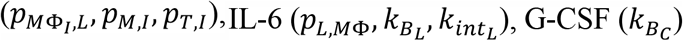, and IFN 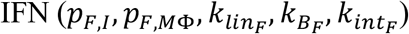.

**Figure 5.**
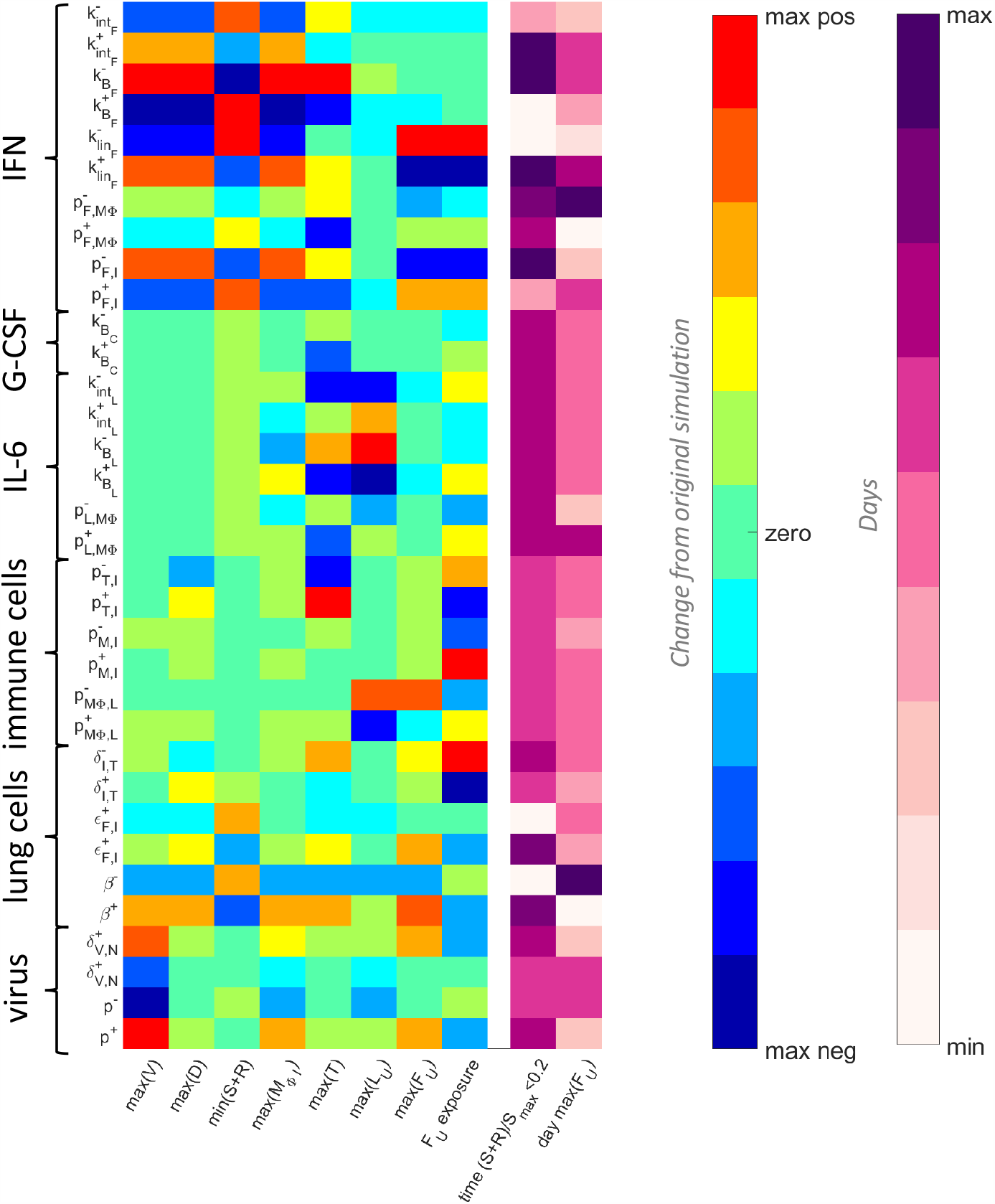
Parameters driving COVID-19 severity. A local sensitivity analysis was performed by varying each parameter ±20% from its originally estimated value and simulating the model. Predictions were then compared to baseline considering: Maximum viral load (max(*V*)), maximum concentration of dead cells (max(*D*)), minimum uninfected live cells (min(*S+R*)), maximum concentration of inflammatory macrophages (max(*M*_*ΦI*_)), maximum number of CD8^+^ T cells (max(*T*)), maximum concentration of IL-6 (max(*L*_*U*_)), maximum concentration of type I IFN (max(*F*_*U*_)), the total exposure to type I IFN (*F*_*U*_ exposure), the number of days damaged tissue was >80% (time (*S* + R)/S_*max*_), and the day type I IFN reached its maximum (day max(*F*_*U*_)). The heatmaps show the magnitude change of each metric, where blue signifies the minimum value observed and red signifies the maximum value observed, or by the number of days, where light to dark pink signifying increasing number of days from zero. The most sensitive parameters are shown here (for complete parameter sensitivity results, see **Figure S8**).

The rate of viral infectivity (*β*) had a particularly significant impact on all metrics where increases resulted in higher viral loads and longer periods of tissue damage > 80%. The duration of extensive tissue damage (>80% damaged) also increased with IFN potency (*ϵ*_*F,I*_). Decreasing the rate of IL-6-induced monocyte differentiation into inflammatory macrophages (*p*_*MΦ,L*_) increased the peak of both IL-6 and IFN. Notably, changes to parameters that increased the bound IFN concentration, i.e. increasing the binding and production rates (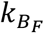 and *p*_*F,I*_) and decreasing the internalization and clearance rates (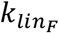 and 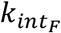) induced significant changes in most metrics (**Figure 5**). See **Figure S8** for complete sensitivity analysis results.

### Virtual patient cohort identifies heterogeneity in immune dynamics and severity

To better understand the clinical variability in SARS-CoV-2 infection severity [1], we next generated a cohort of 200 virtual patients (see **Materials and Methods** and **Figure 7**). To create each *in silico* patient, seven patient-specific parameters were sampled from normal distributions with means corresponding to their respective fixed values and standard deviations inferred from clinical observations (**Table 1**). In doing this, we assumed intrinsic interindividual heterogeneity in monocyte to macrophage differentiation, production of IL-6 by macrophages, recruitment of macrophages by the presence of infected cells, and production of IFN by infected cells, macrophages and monocytes, respectively.

**Table 1.** Virtual patient-specific parameter values. Seven parameters in the model were deemed patient-specific and were drawn from a normal distribution with mean the parameter value obtained either through fitting or from the literature (**Table S1**). The standard deviation (Std Dev) for each normal distribution was informed by values in the literature (see **Materials and Methods** and Supplementary Information Sections S6.1). Initial parameter sampling and new parameters generated through the simulated annealing optimization, were bounded within the interval range noted. All other parameters in the model were fixed to their original value (**Table S1**).

| Parameter | Units | Description | Mean | Ref | Std Dev | Ref | Range |
| --- | --- | --- | --- | --- | --- | --- | --- |
| $p_{M\Phi,I,L}$ | 1/day | Monocyte to macrophage differentiation by IL-6 | 1.7 | [46] | 2.2 | [7] | [0, 9.9] |
| $p_{L,M\Phi}$ | pg/ml/day | IL-6 production by activated macrophages | 1872 | [47] | 2.2 | [7] | [1863, 1880] |
| $p_{F,I}$ | pg/ml/day | IFN production by infected cells | 2.82 | [48] | 1.9 | [44] | [0, 12.2] |
| $p_{M,I}$ | 1/day | Monocyte recruitment rate by infected cells | 0.22 | [49] | 0.08 | [50] | [0, 0.63] |
| $\eta_{F,M\Phi}$ | $10^9$cells/ml | IFN by infected cells | 0.0012 | [48] | $10^{-5}$ | [51] | [0, $10^{-4}$] |
| $\epsilon_{F,I}$ | pg/ml | IFN production of CD8<sup>+</sup> T cells | 0.004 | [52] | $10^{-5}$ | [45] | [0, $10^{-4}$] |
| $p_{F,M}$ | pg/ml/day | IFN production by monocytes | 3.56 | [53, 54] | 0.013 | [53] | [3.4, 3.6] |

Parameters were chosen based on their impact on maximum IL-6 and IFN levels as well as tissue damage observed in the sensitivity analysis (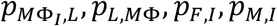, and *ϵ*_*F,I*_; **Figure 5**). In addition, we designated patient-specific parameters accounting for alternate pathways through which IFN is affected by innate immune cells (*η*_*F*, *MΦ*_and *p*_*F,M*_). For the production of IL-6 by macrophages and monocyte to macrophage differentiation via IL-6 stimulation, standard deviations were inferred from IL-6 levels in non-mechanically ventilated patients (mild) and from mechanically ventilated patients (severe) [44] (**Figure S7**D). Standard deviations for the production of IFN by infected cells were determined from the 95% confidence interval for IFN-*α* from Trouillet-Assant et al. [42] (**Figure S7**A-B), and, lastly, the standard deviation for the production of IFN by macrophages was obtained from the 95% confidence interval in Sheahan et al. [45]. The variation in virtual patient responses was then constrained by experimental and clinical viral loads, IFN, neutrophil, IL-6, and G-CSF (**Figure 7**). The resulting cohort dynamics were within ranges for IFN and IL-6 measurements in asymptomatic to severe COVID-19 patients in the literature [11, 17] (**Figure S9**).

To quantify disease severity, we introduced an inflammation variable, Ψ, that measured maximum IL-6, neutrophils and tissue damage (**Eq. 18**) and then compared it to individual characteristics of each virtual patient’s disease. We evaluated each virtual patient’s maximum IL-6, CD8^+^ T cells, and neutrophils; minimum percentage of healthy lung tissue; the time to peak IFN; and total IFN exposure (area under the curve or AUC) within 21 days of infection. Ordering patients by their value of Ψ and plotting the corresponding values for different characteristics evaluated showed a clear separation between those with mild disease and those with severe disease (**Figure 6A**).

**Figure 6.**
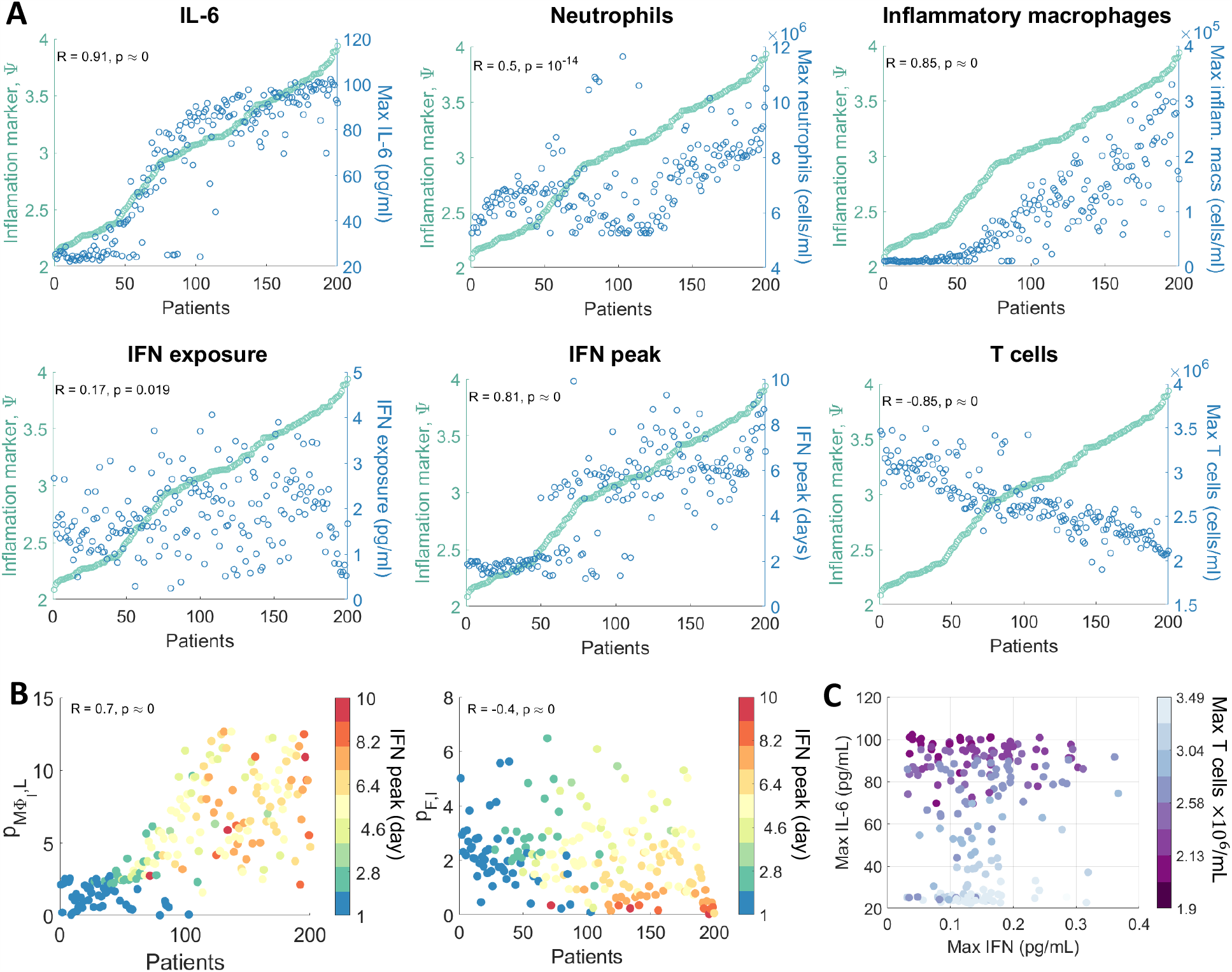
Virtual Cohort of SARS-CoV-2 infected patients. 200 virtual patients were generated by sampling parameters related to macrophage, IL-6, and IFN production (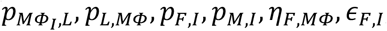 and *p*_*F,M*_) from normal distributions with mean equal to their original values and standard deviation inferred from clinical observations (**Figure 7)**. Each virtual patient had a distinct parameter set that was optimized to that patient’s dynamics in response to SARS-CoV-2 infection corresponded to physiological intervals reported in the literature (see **Materials and Methods**). **A)** Infection and immune response metrics (blue) in individual patients were compared to inflammatory variable Ψ (green). Each point represents an individual patient, ordered according to Ψ. The correlation coefficient (R) and p-value are indicated for each, with α<0.05 denoting significant correlations. **B)** Parameters most correlated to the IFN peak time were the rates of macrophage production via IL-6 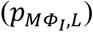 and the IFN production by infected cells (*p*_*F,I*_). Individual patient values for these parameters are plotted as circles coloured by the patient’s corresponding day of IFN peak (see color bar). Patients are ordered by their inflammation marker Ψ. **C)** Correlations between maximal IFN, IL-6, and T cell concentrations for each patient (circles). Circle colour corresponds to the maximal T cell concentration of each patient.

Patients with higher inflammation had higher IL-6, neutrophil, and inflammatory macrophage concentrations (**Figure 6A**). While the IFN exposure was not significantly stratified by Ψ, the peak of IFN and CD8^+^ T cell levels were strongly negatively correlated with the inflammation marker (R = −0.85, p < 1 × 10^−9^, see **Materials and Methods**). IL-6 was most noticeably correlated with Ψ (R = 0.91, p< 1 × 10^−9^), with a distinct upper bound in the concentration (∼100 pg/ml) achieved in 50% of the virtual cohort. There appeared to be a transition phase in inflammation driven by neutrophil levels where patients with moderate inflammation (3 < Ψ < 3.5) had low counts (less than 7 × 10^6^ cells/ml) compared to patients with more severe inflammation (Ψ ≥ 3.5) who had higher levels (p = 1.46 × 10 ^−6^). Despite this, patients with moderate inflammation exhibited increased disease markers including delayed IFN peaks and lower CD8^+^ T cells, compared to patients with mild inflammation (Ψ ≤ 3).

A distinct jump in the timing of the IFN peak in the virtual cohort (p < 1 × 10^−5^) was correlated with inflammation, as patients with low inflammation (Ψ ≤ 3) had peaks at day 2 compared to day 6 in patients with higher inflammation (Ψ>3). Grouping virtual individuals by their time to IFN peak suggests that those with IFN peaks after day 3 of infection also had fewer macrophages (p< 1 × 10^−5^) and larger numbers of CD8^+^ T cells (p < 1 × 10^−5^). Overall, delays in IFN peak did not cause significant changes to viral load but were sufficient to cause major tissue damage (100x reduction in viable tissue remaining) and over-heightened immune responses (4x increase in maximum IL-6 and GM-CSF concentrations).

We found a positive correlation (R= 0.67, p= 1.58 × 10^−8^) between the time to peak IFN concentration for each patient against their IFN production rate from infected cells (**Figure 6B**). Interestingly, the time to peak IFN for each patient was also strongly related to their rate of IL-6-stimulated monocyte differentiation into macrophages. Low IFN production rates were predominately responsible for significantly delayed IFN peaks over 6 days after infection, whereas IFN peaks within 3 days of infection were largely caused by lower rates of monocyte to macrophage differentiation.

Further, examining the relationship between each virtual patient’s maximum IL-6, IFN, and CD8^+^ T cell concentrations (**Figure 6C**) identified a weaker correlation between the maximum concentration of CD8^+^ T cells and IFN (R= 0.24, p = 0.0008) as opposed to with IL-6 (R= −0.86, p < 1 × 10^−9^).

## Discussion

Serial immunological measurements from COVID-19 patients are only beginning to be collected, and the ability to assess initial infection kinetics and the drivers of the ensuing disease remains limited. The data-driven mechanistic mathematical model and virtual patient cohort developed here identified important immunological drivers of COVID-19. In particular, to recreate severe dynamics, it was sufficient to vary only two processes in the model: the rates of type I IFN production from infected cells and macrophages, and the rate of monocyte recruitment by infected cells. This suggests that the distinction between severe and mild disease may be driven by a limited set of causal regulators. The effect on IFN production may be further exacerbated by autoimmunity against type I IFNs, which has been shown to correlate to life-threatening COVID-19 pneumonia in 2.6% of women and 12.5% of men [18].

Our results show that delaying type I IFN production is sufficient to cause major tissue damage and heightened immune responses yet have little impact on peak viral loads. In the severe disease simulation, viral load was cleared marginally faster (∼1 day) in comparison to the mild disease simulation. This finding is supported by recent clinical evidence showing that an increased rate of viral decline rather than peak viral load may be more predictive of disease severity [6]. This therefore suggests that viral load may not be a necessary attribute to obtain severe tissue damage. Instead, our model predicts that increases in tissue damage occur through heightened innate immune responses. Evaluating SARS-CoV-2 infection in a cohort of 200 virtual patients revealed several immunological responses responsible for differential disease presentation. Notably, a distinct, emergent switch in the type I IFN response corresponded with late IFN peaks and more severe disease (i.e., higher inflammation *Ψ*). This supports previous findings that connect a delay in type I IFN with more severe presentations of highly pathogenic coronaviruses infections including SARS-CoV, MERS-CoV, and SARS-CoV-2 [13, 14, 22]. Virtual patients with rates of monocyte differentiation close to the rate at homeostasis tended to achieve peak IFN concentrations approximately 2 days after infection compared to those with higher inflammation and later IFN responses, who had at least a 3-fold increase in this rate. This switch in timing was caused by increased rates of monocyte-to-macrophage differentiation and decreased production of IFN by infected cells, with the initial delay of IFN caused by increased monocyte differentiation and the more extreme IFN delays caused by IFN production from infected cells, indicating that the timing of the IFN peak in a patient may allow for improved stratification into treatment arms designed to target one or both of these responses. The finding that IFN binding was predictive of the duration of lung tissue damage, suggests that virus-intrinsic properties and their ability to inhibit receptor mediated binding and endocytosis could delay IFN production and cause downstream increases in IL-6 and GM-CSF resulting in severe disease. Our results further highlight that lymphopenia is tightly correlated with maximum IL-6 concentration and less dependent on the timing of IFN.

The ability of our model to recapitulate severe disease by, in part, regulating monocyte differentiation raises the possibility that patients with low monocyte levels [7] may benefit from treatments that better regulate monocyte differentiation. This is in line with recent studies identifying distinct transcriptional factors as regulators of differentiated monocyte fates in inflammatory conditions [55, 56]. It also raises the possibility that modulation by exogenous cytokines, including macrophage colony-stimulating factor in combination with IL-4 and tumour necrosis factor-alpha (TNF-*α*), may be able to direct monocyte differentiation in favour of monocyte-derived dendritic cells and reduce this response [55]. Recently, the neutralization of both TNF-α and IFN-γ has been found to benefit patients with COVID-19 or other cytokine storm-drive syndromes by limiting inflammation and tissue damage [57]. Given that TNF-α also has a secondary benefit on monocyte differentiation, our results support the viability of this avenue of treatment. Caution should be noted, however, given that previous attempts to regulate host responses by IL-6 blockade have proven unsuccessful [58].

Together, our findings support the idea that early interventions aimed at reducing inflammation are more likely to be beneficial for patients at risk of progressing to severe COVID-19 than attempts to inhibit cytokine storm later in the disease course, given that early IFN responses were found to provoke better controlled immune responses and outcomes in our virtual cohort. It will be essential to characterize both the timing and mechanisms of proposed therapeutic interventions to develop effective treatments to mitigate severe disease.

## Materials and Methods

### Mathematical model of the immune response to SARS-CoV-2

Our model was developed to examine SARS-CoV-2 infection dynamics and identify immunological drivers of disease severity (**Eqs. S1-S22**). Throughout, cytokine and immune cell interactions and effects were described by Hill functions as

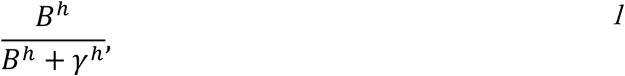

where *B* is the interacting compound, *γ* its half-effect value, and h the Hill coefficient [59, 60]. Further, for a given cytokine *X* and cell population *Y*, the production (recruitment/differentiation) rate of *X* by *Y* was denoted by *p*_*Y,X*_ and the rate of production of *Y* by *X* by *p*_*X,Y*_. The half-effect concentration (i.e. *γ* in **Eq. 1**) of cytokine *X* on cell population *Y* was represented by *ϵ*_P,*X*_ and the half-effect concentration of cell *Y* affecting cytokine *X* was given by *η*_X,*Y*_. The natural death rate of cell *Y* was denoted by *d*_*Y*_, and the rate of induced death of cell *Y* by cell *Z* by *δ*_*Y,Z*_. Lastly, the carrying capacity concentration of cell *Y* was denoted by *Y*_*max*_, and regeneration or proliferation rates by *λ*_*Y*_.

We modelled virus (*V*) being produced by infected cells at rate *p* and cleared via exponential clearance at rate *d*_*V*_, which accounts for all contributions to viral degradation except macrophage- and neutrophil-mediated clearance. Immune-mediated viral clearance via phagocytosis by inflammatory macrophages [61] and neutrophil extracellular traps (NETs—extracellular chromatin fibres produced by neutrophils to control infections) [39, 40] was considered to occur at rates 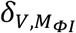 and *δ*_*V,N*_, respectively. Susceptible epithelial cells (*S*) grow logistically with per capita proliferation rate *λ*_*S*_ and carrying capacity *S*_*max*_, and become infected (*I*) at rate *β*. The damage inflicted on epithelial cells by neutrophils was modelled using a Hill function (**Eq. 1**) [60], where neutrophils kill/damage epithelial cells at rate *δ*_*N*_ through the release of NETs and other antimicrobials proteins [39, 40]. The constant *ρ* (0 < *ρ* < 1) was included to modulate bystander damage of uninfected cells (*S* and *R*).

For the purposes of our investigation, we only considered type I IFN dynamics (primarily IFN-α, β). Type I IFN (*F*_*U*_ and *F*_*B*_) reduces the infectivity and replication capability of viruses by stimulating cells to become resistant to infection. These resistant cells (*R*) proliferate at a rate equivalent to susceptible cells (*λ*_*S*_). The concentration of bound IFN (*F*_*B*_) modulates the creation of infected and resistant cells [19, 21, 62, 63], where increasing the concentration of IFN causes more cells to become resistant to infection and less to become productively infected (*I*). The potency of this effect is controlled by the half-effect parameter *ϵ*_*F,I*_. Following the eclipse phase (which lasts *τ*_I_ hours), productively infected cells (*I*) produce virus before undergoing virus-mediated lysis at rate *d*_*I*_. Although various immune cell subsets contribute to infected cell clearance, we limited our investigation to macrophages and effector CD8^+^ T cells which induce apoptosis at rates *δ*_*I*,.*M Φ*_and *δ*_I,*T*_, respectively.

The accumulation of dead cells (*D*) was assumed to occur through infected cell lysis *d*_*I*_, neutrophil damage/killing of epithelial cells *δ*_*N*_, macrophage phagocytosis of infected cells *δ*_*I,MΦ*_, macrophage exhaustion *δ*_.*MΦ*,D_, and CD8^+^ T cell killing of infected cells *δ*_*I,T*_. These dead cells disintegrate relatively quickly [64] at rate *d*_*B*_, and are cleared through phagocytosis by macrophages [65] at rate *δ*_*D*,.*MΦ*_.

Resident alveolar macrophages (*M*_*ΦR*_) are replenished at a logistic rate inversely proportion to viral load with maximal rate of *λ*_.MΦ_ and half-effect *ϵ*_*V*,.*MΦ*_ (i.e. as the virus is cleared, the inflammatory macrophage pool replenishes the alveolar macrophage population in the lung). We modelled the transition of alveolar macrophages to inflammatory macrophages (*M*_*ΦI*_) as dependent on infected and dead cells, with a maximal rate of *a*_*I,.MΦ*_. Resident macrophages die naturally at a rate 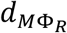 or due to the clearing of dead cells (exhaustion) [65] at rate *δ*_.*MΦ,D*_.

Inflammatory macrophages are produced by three distinct pathways (acting individually or in concert): 1) stimulated tissue-resident macrophages *a*_*I.M*Φ_, (2) GM-CSF-dependent monocyte differentiation, with maximal production *p*_*M*,G_ and half effect *ϵ*_*G,M*_, and (3) IL-6-dependent monocyte differentiation, with maximal production rate 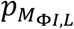 and half-effect 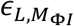. We assumed that inflammatory macrophages die naturally at rate 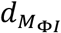 or from clearing dead cells at a rate *δ*_.*MΦ,D*_.

We have previously shown that endogenous cytokine concentrations are far from quasi-equilibrium at homeostasis [66]. Therefore, to describe the pharmacokinetics and pharmacodynamics of cytokine binding and unbinding, we leveraged the framework established in Craig et al. [66] (**Figure 1C**) for IFN (*F*_B_ and F_U_), IL-6 (*L*_*B*_ and *L*_*U*_), GM-CSF (*G*_*B*_ and *G*_*U*_), and G-CSF (*C*_*B*_ and *C*_*U*_). In its general form, this pharmacokinetic relationship is expressed as

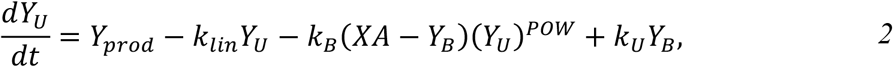

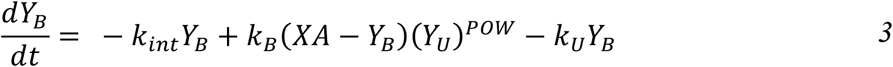

where *Y*_*U*_ and *Y*_*B*_ are free and bound cytokines, *Y*_*prod*_ is the rate of endogenous cytokine production, *k*_*B*_ and *k*_*U*_ are the respective binding and unbinding rates, *k*_*int*_ is the internalization rate of bound cytokine, and *k*_LiN_ is the elimination rate. Here, *POW* is a stoichiometric constant, *A* is a scaling factor and *X* is the sum of all cells modulated by the cytokine with

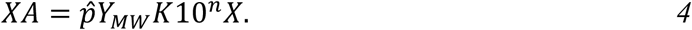

where 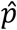 is a constant relating the stoichiometry between cytokine molecules and their receptors, *K* is the number of receptors specific to each cytokine on a cell’s surface and 10^*n*^ is a factor correcting for cellular units (see **Eqs. S19-S22**). The molecular weight was calculated in the standard way by dividing the cytokine’s molar mass (*MM*) by Avogadro’s number (*Y*_*MW*_ = MM/6.02214 × 10^23^).

We considered unbound IL-6 (*L*_U_) to be produced from productively infected cells, inflammatory macrophages, and monocytes, with bound IL-6 (*L*_*B*_) resulting from binding to receptors on the surface of neutrophils, CD8^+^ T cells and monocytes. Unbound GM-CSF (*G*_*U*_) was assumed to be produced from inflammatory macrophages and monocytes and bind to receptors on monocytes to create bound GM-CSF (*G*_B_). GM-CSF can be produced by CD8^+^ T cells [67], but this was excluded because it was insignificant to the full system’s dynamics. Unbound G-CSF (*C*_*U*_) is secreted by monocytes, with bound G-CSF (*C*_B_) produced via binding to neutrophil receptors. Lastly, because unbound type I IFNs (*F*_*U*_) are known to be produced by multiple cell types in response to viral infection, including lymphocytes, macrophages, endothelial cells and fibroblasts [62], we modelled its unbound production from infected cells, infiltrating/inflammatory macrophages, and monocytes, and its binding to receptors on both CD8^+^ T cells and infected cells (**Figure 1B**).

The pharmacokinetics and pharmacodynamics of G-CSF on neutrophils (*N*) were taken directly from Craig et al. [66]:

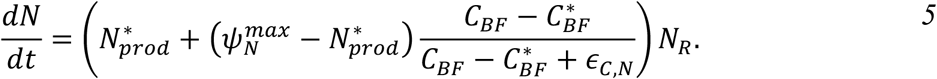

Neutrophil recruitment of bone marrow reservoir neutrophils (*N*_*R*_) was modelled to occur via the bound fraction of G-CSF [68] (*C*_*B*F_ *= C*_*B*_*(t)/(A*_*C*_*N(t)*)) at rate 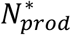 which increases towards its maximal value 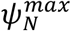 as a function of increasing G-CSF. During the acute phase of inflammation, endothelial cells produce IL-6 leading to the attraction of neutrophils [69]. This was modelled as recruitment with maximal rate *p*_*N,L*_ and half-effect parameter *ϵ*_*D,L*_. Neutrophils die at rate *d*_*N*_.

Monocytes (*M*) are recruited by bound GM-CSF [70], similar to neutrophils (**Eq. 5**), with bone marrow monocytes (*M*_*R*_) recruited at a homeostatic rate 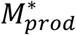. In the presence of GM-CSF, this rate increases towards 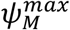. Monocytes are also recruited by the presence of infected cells at a maximal rate of *p*_*M,I*_ with half-effect *ϵ*_*I,M*_, and subsequently disappear through differentiation into inflammatory macrophages (as above) or death at rate *d*_*M*_.

CD8^+^ T cells are recruited through antigen presentation on infected cells as a function of infected cell numbers at rate *p*_*T,I*_ The constant delay (*τ*_*T*_) accounts for the time taken for dendritic cells to activate, migrate to the lymph nodes, activate CD8^+^ T cells, and the arrival of effector CD8^+^ T cells at the infection site. CD8^+^ T cell expansion occurs in response to bound IFN at a maximal rate *p*_*T,F*_ with half-effect *ϵ*_*F,T*_, and CD8^+^ T-cell exhaustion occurs with high concentrations of IL-6 [16, 17], with half-effect *ϵ*_*L,T*_, and apoptosis occurs at rate *d*_*T*_. All variable and parameter descriptions are provided in **Table S1**.

### Estimating early infection dynamics (‘viral model’)

To begin estimating parameter values from data, we set all immune populations and cytokine concentrations in the full model (Supplementary Information **Eqs. S1-S22)** to zero (*M*_Φ*R*_ *= M*_Φ*I*_ *= M = N = T = L*_*U*_ *= L*_*B*_ *= G*_*U*_ *= G*_*B*_ *= C*_*U*_ *= C*_*B*_ *= F*_*U*_ *= F*_*B*_ *= 0*). This gives

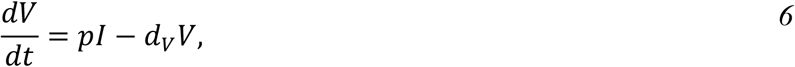

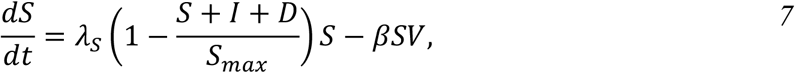

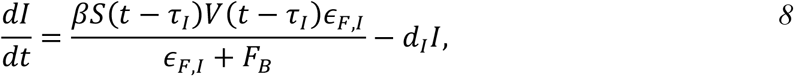

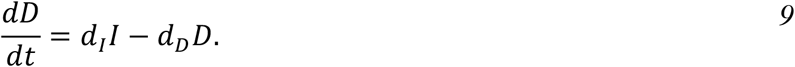

We also assumed there were no resistant cells (*R = 0*) due to the absence of an IFN equation. This resulted in a simplified ‘viral model’ that considers only virus (*V*) infection of susceptible cells (*S*) which creates infected cells (*I*) after *τ*_*I*_ days, which the die through lysis, creating dead cells (*D*).

### Type I interferon dynamics during early infection (‘IFN model’)

To study infection dynamics driven uniquely by IFN, we extended **Eqs. 6**-**9** by introducing the IFN mechanisms from **Eqs. S1-S22**, i.e. setting other cytokine and immune cell populations to zero (*M*_Φ*R*_ *= M*_Φ*I*_ *= M = N = T = L*_*U*_ *= L*_*B*_ *= G*_*U*_ *= G*_*B*_ *= C*_*U*_ *= C*_*B*_ *= 0*), giving

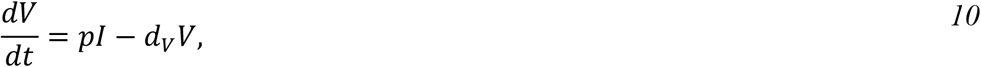

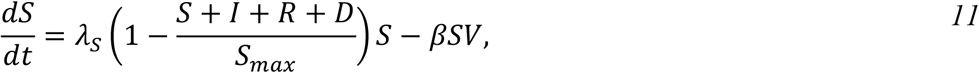

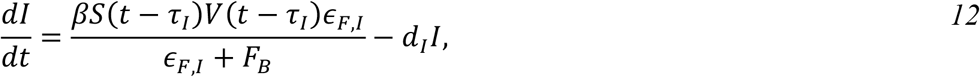

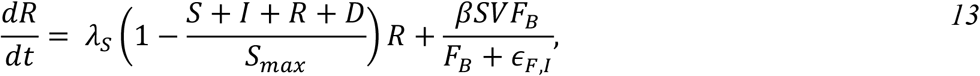

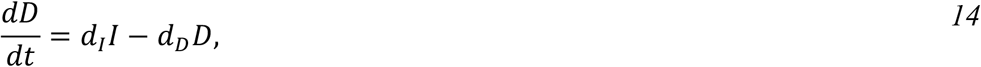

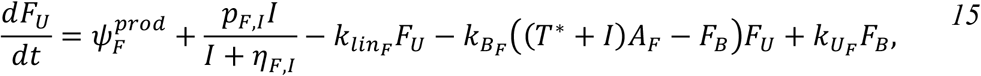

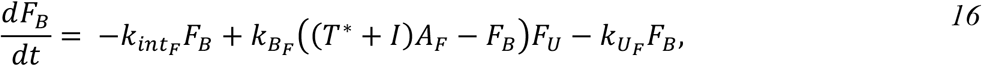

where cells become resistant (*R*) through IFN (*F*_*U*_ and *F*_*B*_). The parameter 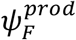 was introduced to account for the production of IFN by macrophages and monocytes not explicitly modelled in this reduced system but included in the full system (i.e. *p*_*F,M*_ and *p*_*F,M*Φ_ in **Eq. S17**). Previously-fit parameters were then fixed to their estimated values (**Table S1**) and the value of 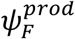 was determined by solving *dF*_*U*_*/dt = 0* at homeostasis (i.e. *V = I = 0*), giving 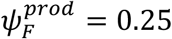.

### Model calibration and parameter estimation

Model parameters (**Table S1**) were obtained either directly from the literature, through fitting effect curves (**Eqs. S24-S25**) or sub-models (**Eqs. S26-S56**) to *in vitro* or *in vivo* data, or by calculating the value that ensured that homeostasis was maintained (**Eqs. S57-S70**) in the absence of infection. All fitting procedures were performed using MATLAB 2019b functions *fmincon* or *lsqnonlin* [71].

Initial concentrations of all unbound cytokines (*L*_*U*,0_, *G*_*U*,0_, *C*_*U*,0_ and *F*_*U*,0_), susceptible cells, resident macrophages, monocytes, neutrophils, and CD8^+^ T cells (*S*_0_, *M*_Φ*R*,0_, *M*_0_, *N*_0_ and *T*_0_) were estimated from plasma and lung tissue concentrations in humans. Parameters for cytokine binding and unbinding kinetics (**Eqs. 2-4**), such as the molecular weight (*MM*), binding sites per cell (*K*), binding/unbinding rates (*k*_*B*_ and *k*_*U*_), internalization rates for GM-CSF, G-CSF and IFN (*k*_*234*_), and cytokine clearance rates (*k*_*lin*_), were estimated both from experimental measurements and previous modelling work. The stoichiometric constants *POW* and 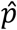 were both equal to 1 for all cytokines, except for G-CSF for which *POW = 1*.*4608* and 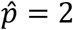 as previously estimated by Craig et al. [66]. Neutrophil and monocyte reservoir dynamics, monocyte differentiation, macrophage activation, and CD8^+^ T cell recruitment and expansion parameters were primarily estimated from previous mathematical modelling studies. Immune cell death rates were taken directly from the literature or estimated from recorded half-lives using **Eq. S23**.

The rates of virus production, decay, infectivity, and infected cell lysis (*p, d*_*V*_, *β* and *d*_*I*_ respectively) were then estimated by fitting **Eqs. 6-9** to viral load measurements from SARS-CoV-2 infection in macaques [41] where eight adult rhesus macaques inoculated with *4 × 10*^*7*^ TCID_50_/ml (*3 × 10*^*’*^ genome copies/ml) SARS-CoV-2 [41] (**Table S1**). Viral loads below 1 copy/ml were assumed to be negligible. Estimated parameters for viral decay and cell lysis (*d*_*V*_ and *d*_*I*_) were used as an upper bound for parameter values in the full model.

A subset of parameters was obtained through fitting sigmoidal effect curves (**Eqs. S24-S25**) curves to *in vitro* and *in vivo* experiments. These include the half-effect neutrophil concentration for epithelial cell damage, the half-effect concentrations for monocyte production and differentiation through GM-CSF signalling (*ϵ*_*G,M*_ and 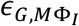 **Figure S1**). Other parameters obtained through effect curves were the half-effects for IL-6 production by monocytes (*η*_*L,M*_), the effect of IL-6 on monocyte differentiation (*ϵ*_*L,M*_) and the half-effect of IFN on CD8^+^ T cell (*ϵ*_,,*T*_) and IL-6 on CD8^+^ T cell expansion (*ϵ*_*L,T*_) (**Figure S2**).

These parameters were then fixed, and remaining parameters were estimated by fitting time-dependent sub-models of **Eqs. S1-S22** to relevant data. The proliferation rate of epithelial cells (*λ*_*@*_), the internalization rate of IL-6 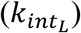, and the rate of neutrophil induced damage were fit to corresponding time-series measurements using exponential rate terms (**Figure S2**). Clearance and phagocytosis of infected cells and extracellular virus by inflammatory macrophages (*δ*_*I,M*Φ_ and *δ*_*V,M*Φ_) were fit to *in vitro* experiments (**Figure S2**). Production of IFN by macrophages (*p*_*F,M* Φ_) was obtained by fitting to data measuring IFN-αproduction (**Figure S3**). The parameters regulating the rate of the resident macrophage pool replenishment (*λ*_*M*Φ_ and *∈* _*V,M*, Φ_) were estimated from *in vivo* observations of resident macrophages during influenza virus infection (**Figure S3**). GM-CSF production by monocytes (*p*_*G,M*_; Figure S3), IFN production by infected cells (*p*_*F,I*_), and IL-6 production by infected cells and macrophages (*p*_*L,I*_ and *p*_*L,M*Φ_) were all obtained from fitting reduced versions of **Eqs. S1-S22** to *in vitro* experiments [47, 48, 72, 73] (**Figure S4**).

Lastly, any remaining parameters values were obtained by ensuring that homeostasis was maintained in absence of infection (**Figure S5**). Parameters calculated from homeostasis include the half-effect monocyte concentration for G-CSF production (*η*_*C*_,_*M*,_), the production rate of IL-6 and GM-CSF by inflammatory macrophages (*p*_*L,M*Φ_ and *p*_*G,M*Φ_), the production rate of monocytes by GM-CSF (*p*_*M,G*_), and the half-effect inflammatory macrophage concentration for IFN production (*η*_*F,M*Φ_). For some parameters it was not possible to obtain an estimation from the literature, and for these we either set their value equal to an already estimated parameter 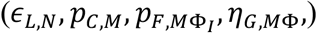, or qualitatively estimated it (*∈*_*I,M*_,*ρ*).

For the ‘IFN model’ (**Eqs. 10-16**), parameters related to virus (*p, d*_*V*_, *β* and *d*_*I*_), epithelial cell proliferation (*λ*_*S*_ and *S*_*max*_), and IFN (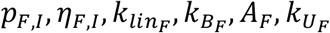 and *∈*_,,*$*_) were fixed to those in **Table S1**.

### Numerical simulations

All ODE models were solved using *ode45* in MATLAB, and delay differentiation equations (i.e. **Eqs. S1-S22**) were solved using *ddesd* in MATLAB.

### Sensitivity analysis

We performed a local sensitivity analysis for the full model (**Eqs. S1-S22)** by individually varying each parameter by ±20% from its estimated value and quantifying the effect on the model’s output. This change was recorded and used to evaluate different metrics representing the inflammatory response to SARS-CoV-2, namely maximum viral load, maximum number of dead cells, minimum uninfected tissue, maximum number of inflammatory macrophages, maximum number of CD8^+^ T cells, maximum unbound IL-6, maximum unbound IFN, the total exposure (AUC) to type I IFN, number of days the percent of damaged tissue was >80%, and time of unbound type I IFN peak. We quantified the fraction of undamaged tissue by (*S* + *R*)/*S*_*max*_.

### Virtual patient generation

To generate a cohort of 200 virtual patients, we followed techniques similar to those of Allen et al. [26] and our previous studies [74, 75] wherein individual virtual patients were created by sampling a parameter set ***p*** from parameter distributions then simulating the model to verify that each individual’s trajectory was realistic. A subset of parameters (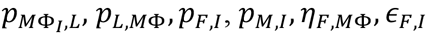, and *p*_*F,M*_) was designated as patient-specific after considering the results of the sensitivity analysis and standard deviations inferred from clinical observations (Supplementary Information). To avoid the inclusion of unrealistic dynamics, patient parameter sets were then optimized using simulated annealing to ensure predictions fell within physiological ranges for viral load [41], IL-6 [6, 44], IFN-α[42], and G-CSF [24] (**Figure 7**).

**Figure 7.**
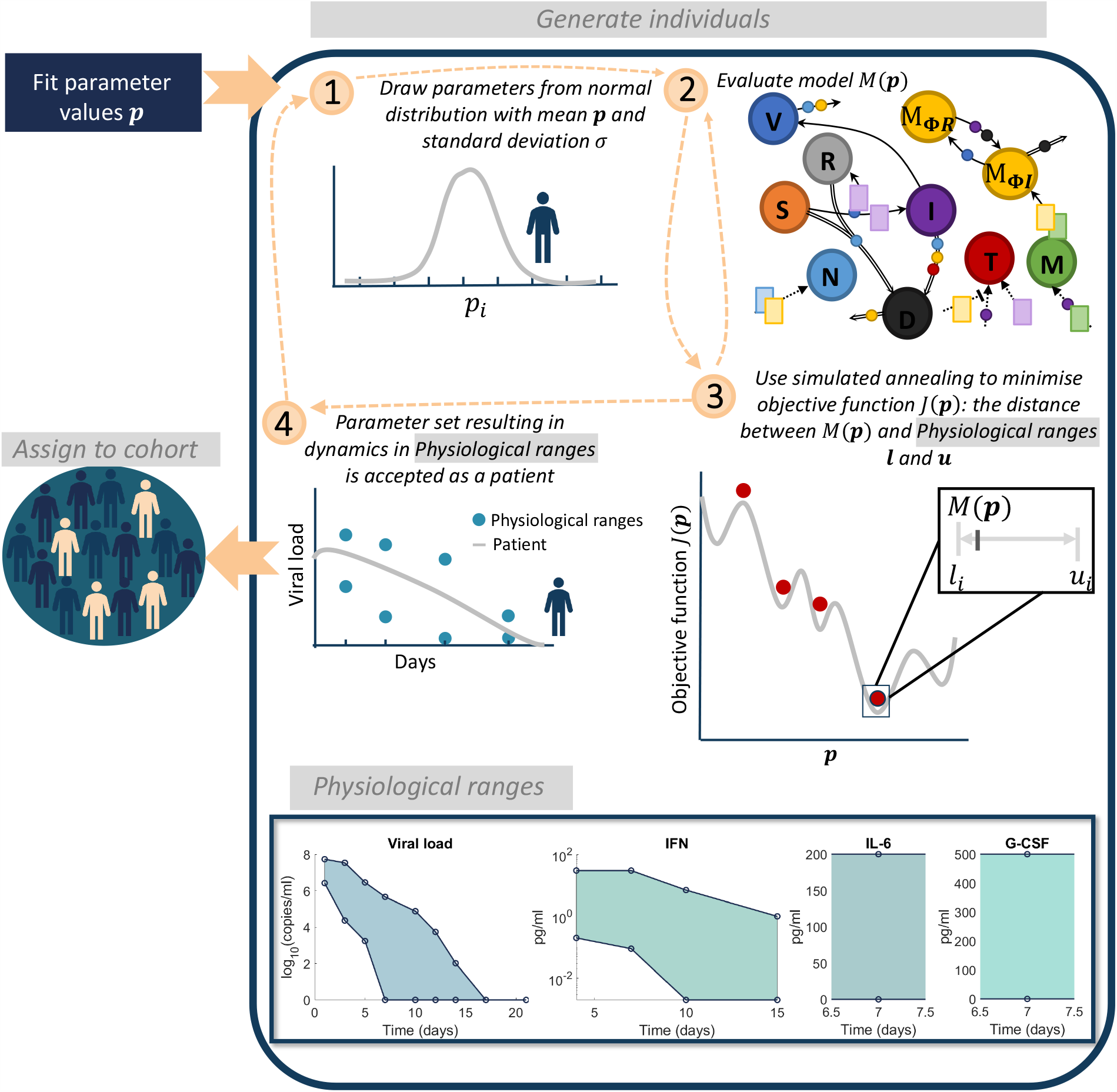
Algorithm for generating virtual patients. Parameters in the model were first obtained through fitting to data (**Table S1**). **1)** Parameters relating to macrophage, IL-6 and IFN production (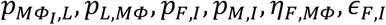, and *p*_*F,M*_) were generated from normal distributions with mean equal to their original fitted values and standard deviation informed by experiment observations (see Section S6.1). **2)** The model evaluated is then evaluated on this parameter set to obtain *y*(*t, p*). **3)** A simulated annealing algorithm is then used to determine a parameter set that optimises the objective function *J*(*p*) (**Eq**.**16**). **4)** Optimizing the objective function provides a parameter set for which the patient response to SARS-CoV-2 will be within the physiological ranges. This patient is then assigned to the cohort and this process is continued until 200 patients have been generated. Physiological ranges are noted in the bottom box for viral load [41], IFN [42], IL-6 [44] and G-CSF [7].

The upper *u*_*I*_ and lower *l*_*I*_ bounds for *V, L*_*U*_, *F*_*U*_ and *C*_*U*_ were based off these physiological ranges from Munster et al. [41] (viral loads), Herold et al. [44] (IL-6 concentrations), Trouillet-Assant et al. [42] (IFN dynamics), and Liu et al. [7] (G-CSF concentrations) as described in Supplementary Information Section S.6.1. Intervals for each patient-specific parameter set were restricted to four standard deviations from the mean or zero if the lower bound was negative. Given an initial patient specific parameter set ***p***, we used simulated annealing to minimize *J*(***p***), i.e.

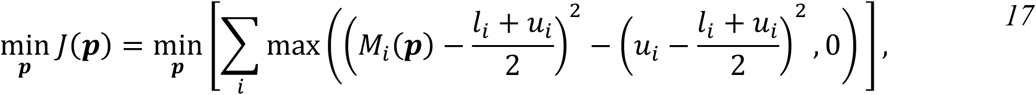

where *M*_*I*_(***p***) is the model output *i* evaluated at parameter set ***p*** corresponding to the upper and lower bound *l*_*I*_ and *u*_*I*_ (**Figure 7**).

To quantify disease severity for each patient, we introduced an inflammation variable (Ψ) to account for the combined changes in IL-6 (*L*_*U*_), neutrophils (*N*), and damaged tissue (*S* + *R*), each normalized by the virtual cohort’s average. In this way, Ψ measures an individual’s relative change from the cohort’s baseline, and quantifies the contributions of IL-6, neutrophils, and tissue damage on comparable scales. For a given patient *j*, the inflammation marker is given by

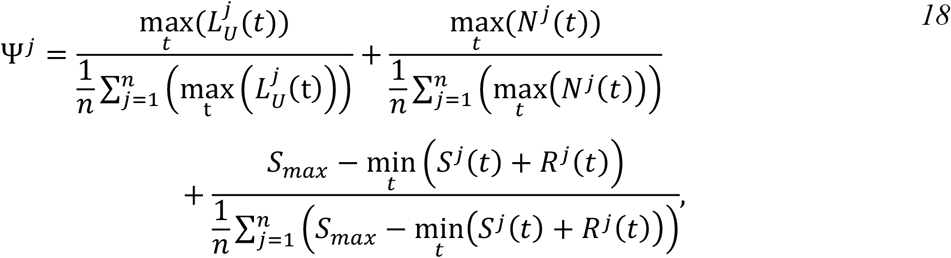

where *n* is the total number of patients in the cohort, and 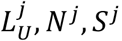, and *R*^*j*^ are the unbound IL-6, neutrophils, and susceptible and resistant epithelial cell count, respectively.

### Statistical analyses

The Pearson correlation coefficient (R) was used to measure the degree of interaction between two variables, with a significance level of *α* < 0 05 indicating rejection of the hypothesis that there is no relationship between the observed variables. In addition, we used two-sample two-sided t-tests (number of patients < 40) and z-tests (number of patients ≥ 40) at the α< 0 05 significance level to test the hypothesis that there were no differences between sample means.

## Acknowledgments

All authors would like to thank Paul Macklin and Thomas Hillen for encouraging collaboration.

## Funding

ALJ was supported by Fonds de recherche Santé Québec Programme de bourse de formation postdoctorale pour les citoyens d’autres pays and a Centre for Applied Mathematics in Bioscience and Medicine (CAMBAM) Postdoctoral fellowship. RA and AMS were supported by NIH R01 AI139088. ALJ and MC were supported by NSERC Discovery Grant RGPIN-2018-04546 and NSERC Alliance COVID-19 ALLRP 554923 – 20.

## Author contributions

ALJ, RA, AMS, CLD, and MC conceived the study. AMS and APS conducted the alveolar macrophage and viral load *in vivo* experiments. PAM contributed significant immunological insight. All authors contributed intellectual insight into the development of the study. ALJ and MC wrote the code and manuscript with significant contributions from all authors.

## Competing interests

The authors declare no competing interests.

## Data and materials availability

All data needed to evaluate the conclusions in the paper are present in the paper and/or the Supplementary Materials.”). Sample code is available upon reasonable request to the corresponding author.

## Supporting information captions

### Supplementary Information file

**Figure S1. Effects of neutrophils on lung epithelial cells, GM-CSF on monocyte production and differentiation, the relationships between monocytes and CD4+ T cells with IL-6, and the influence of IFN on T cell expansion. A)** Using the measurements by Knaapen et al.^22^, the inhibitory effect curve E (**Eq. S25**) was fit to the cell viability of RLE cells under various concentrations of H_2_O_2_. **B)** The stimulatory effect curve E (**Eq. S24**) was fit to the dose response measurements of blood monoculture cells (3 × 10^(^cells/dish) with various concentrations of murine recombinant GM-CSF (IU/ml)^18^. **C)** The stimulatory effect curve E (**Eq. S24**) was fit to measurements for the monocytic myeloid cell count as a function of GM-CSF.^17^ **D) Eq. S27** fit to time course data of IL-6 production from monocytes^38^. **E)** IL-6 stimulation of monocyte differentiation to macrophages modelled by the inhibitory effect curve E (**Eq. S24**) fit to the percentage of CD14+ cells (macrophages) as a function of the number of fibroblasts measured by Chomarat et al.^16^. **F)** Stimulatory effect curve E (**Eq. S24**) for IFN-γ stimulation on CD8+ T cells fit to measurements of the signalling in CD8+T cells for varying doses of IFN-γ^19^. Data (black) is plotted as either circles (**D & E**) or mean and standard deviation error bars (**A-C&F**); solid blue line: corresponding fit.

**Figure S2. Dynamics of IL-6 on T cell expansion, epithelial cell growth, IL-6 internalization, neutrophil-induced damage, and macrophage phagocytosis. A)** Effect curve (**Eq. S24**) for the IL-6 effect on T cell expansion fit to measurements CD4^+^ T cells from dilutions of IL-6 by Holsti and Raulet^21^. **B)** Exponential growth curve fit to the growth of A549 cells^2^ **C)** The internalization rate of IL-6 (**Eq. S30**) fit to the fraction of internalized IL-6^47^. **D)** Exponential decay fit to cell viability after H_2_O_2_ administration^24^. **E)** The macrophage clearance of apoptotic material (**Eqs. S31-S33)** was fit to the percentage of macrophages that had engulfed material over 25 hours^27^. **F)** The phagocytosis rate of extracellular virus by macrophages was obtained by fitting **Eqs. S34-S35** to the uptake of virus by macrophages measured by Rigden et al.^23^. Data (black) is plotted as either circles (**A & F**) or mean and standard deviation error bars (**B-E**); solid blue line: corresponding fit.

**Figure S3. Monocyte expansion and type I IFN production by monocytes, alveolar macrophage replenishment after viral infection, and GM-CSF production by monocytes. A) Eq. S37**fit to time course of proliferation of monocytes in culture^42^. **B)** Fit of **Eqs. S38-S39** to the production of IFN-α by monocytes after 24 hours with RSV as a function of the number of days of pre-culturing (1, 2, 4 or 7)^43^. **C)** Correlation between infectious virus titre and RT-PCR copy number for influenza A and B measured by Laurie et al.^88^ The relative TCID_50_ compared to the RNA copies is plotted for each virus strain and the mean as a black dashed line. **D-E)** Fit of **Eqs. S40-S42** to viral loads^87^ and alveolar macrophages from experimental influenza infections. **F)** The production of GM-CSF from stimulated monocytes was recorded by Lee et al.^40^ Using a simplified version of the full model (**Eqs. S43-S46**), we obtained the production rates for monocytes and GM-CSF. Data (black) is plotted as either circles/stars (**B&F**) or mean and standard deviation error bars (**A**,**D-E**); solid blue line: corresponding fit.

**Figure S4. Production of IFN and IL-6 by infected cells and macrophages. A)** Concentration of IFN-*β* released by alveolar epithelial cells in response to stimulation with influenza virus recorded at 8, 16 and 24 hours^41^. **B-C)** IL-6 production by infected cells in response to **A)** H5NA and **B)** H7N9, measured by Ye et al.^36^ Data (black) is plotted as mean and standard deviation error bars with the corresponding fit (**Eqs. S51-S54**) in solid blue. **D)** IL-6 production by macrophages (**Eq. S56**) in response to stimulation with LPS of varying dosage sizes. Shibata et al. ^37^ measured the production of IL-6 for different dosages of LPS and fitting the production rate to this data to obtain *p*_*L,MΦ*_, *η*_*L,MΦ*_.

**Figure S5. Homeostatic disease-free system regulation. A)** To confirm that parameters in the model represented realistic immunocompetent individuals in the disease-free scenario, **Eqs. S1-S22** were simulated where *V*_0_ = 0 and parameters were given by the homeostasis **Eqs. S57-S70**. The initial concentration of G-CSF was perturbed and compared to simulations of the model at homeostasis. Simulations at homeostasis are represented by solid lines (purple) and perturbed simulations as dashed lines (pink). **B)** The maximum residual between variables and their initial conditions at day 50 was measured to confirm that the system was stable for perturbations in all immune cells and cytokines.

**Figure S6. Model validation against human cytokine measurements during SARS-CoV-2 infection. A)** IFN dynamics of the reduced model (**Figure 3 Main Tex**t) overlaid with patient IFN-*α*2 plasma concentrations from Trouillet-Assant et al.^70^ The solid line (purple) represents the unbound IFN dynamics from the reduced model (**Eqs. 27-33)**. Individual patient IFN-*α*2 measurements are plotted as grey circles. Normal IFN-*α*2 concentration in healthy volunteers are indicated by a grey area. **B-F)** Mild and severe dynamics (**Eqs. S1-S22**) corresponding to simulations in **Figure 4 Main Text** and **Figure S7** overlaid with measurements from the literature with solid lines: mild disease dynamics; dashed lines: severe disease dynamics. **B-C)** Plasma IFN-*α* and IL-6 in COVID-19 critically ill patients (n=26) obtained by Trouillet-Assant et al.^70^ overlaid with mild and severe unbound IFN (*F*_U_(*t*)) and mild and severe unbound IL-6 (*L*_U_(*t*)). **D)** IL-6 levels in patients requiring and not requiring mechanical ventilation obtained by Herold et al.^91^ overlaid with mild and severe unbound IL-6 dynamics. **E-F)** IL-6 and G-CSF plasma concentration obtained by Long et al.^92^ in symptomatic “S” and asymptomatic “AS” COVID-19 patients overlaid with corresponding mild and severe model dynamics.

**Figure S7. Predicting mild and severe COVID-19 dynamics (all model variables)**. Extension of results of mild and severe disease dynamics in **Figure 4 Main Text**. Mild disease (solid lines) dynamics obtained by using baseline parameter estimates (**Tables S1**) while severe disease dynamics (dashed lines) were obtained by decreasing the production rate of type I IFN, p_*F,I*_, and increasing the production of monocytes, p_*M,I*_, and their differentiation to macrophages, η_F,MΦ_. **A)** Lung cells concentrations (susceptible cells S(T), resistant cells *R*(T), infected cells *I*(T), dead cells D(T) and virus *V*(T)). Solid black line with error bars indicates macaque data (see **Fig. 2 Main Text**). **B)** Immune cell concentrations (resident macrophages *M*_*ΦR*_(T), inflammatory macrophages *M*_*ΦI*_(T), monocytes *M*(T), neutrophils *N*(T) and T cells *T*(T)). **C)** Bound and unbound cytokine concentrations (IL-6 unbound *L*_U_(T) and bound *L*_B_(T), GM-CSF unbound G_U_(T) and bound G_B_(T), G-CSF unbound C_U_(T) and bound C_B_(T), type I IFN unbound *F*_U_(T) and bound *F*_B_(T)).

**Figure S8. Full analysis of parameters driving COVID-19 severity**. A local sensitivity analysis was performed by varying each parameter ±20% from its originally estimated value and simulating the model. Predictions were then compared to baseline considering: Maximum viral load (max(*V*)), maximum concentration of dead cells (max(*D*)), minimum uninfected live cells (min(S+R)), maximum concentration of inflammatory macrophages (max(*M*_Φ*I*_)), maximum number of CD8^+^ T cells (max(*T*)), maximum concentration of IL-6 (max(*L*_U_)), maximum concentration of type I IFN (max(*F*_U_)), the total exposure to type I IFN (*F*_U_ exposure), the number of days damaged tissue was >80% (time (*S* + R)/S_*max*_)<0.2), and the day type I IFN reached its maximum (day max(*F*_U_)). The heatmaps show the fold change of each metric, where blue signifies the minimum value observed and red signifies the maximum value observed, or by the number of days, where light to dark pink signifying increasing number of days from zero. The most sensitive parameters are shown in **Figure 5** in the **Main Text**.

**Figure S9. Cohort dynamics within physiological ranges**. Virtual patients were generated so that viral load, IFN and IL-6 concentration were within physiological ranges obtained in the literature. The physiological ranges (denoted by open circles) were obtained from **A)** Munster et al.^96^, **B)** Trouillet-Assant et al. ^70^, and **C)** Herold et al. ^91^. Patient dynamics at discrete time points are plotted as joined green dots.

**Table S1. Parameter values used in the Main Text**. Parameters have been grouped into: (a-e) cell related, (f-k) cytokine related parameters (l) and initial conditions. Relevant references are given estimated parameters. Parameters obtained through fitting to data in the literature have the appropriate figure noted in the Info column. Parameters estimated from homeostasis calculation are denoted by H or qualitatively estimated by E. Parameters whose value was taken from another parameters estimated has that parameter noted. Viral load is reported as virion copies and cells have been noted in 10^8^cells. Time *T* is in days. The final sub-table (m) is a list of the variables in the model.

## Notes

### Competing Interest Statement

The authors have declared no competing interest.

