## Supplementary Information for "COVID-19 virtual patient cohort reveals immune mechanisms driving disease outcomes"

In the Supplementary Information, we detail the full model equations (Section S1), the parameters and their values (Section S2), and how parameters were estimated (Section S3-S5). Parameters in the model were either obtained from the literature (Section S3), through fitting to dose response data (Section S4.1) or through fitting to time-series measurements (Section S4.2). Remaining parameters were estimated through calculating homeostasis (Section S5). Validation against human SARS-CoV-2 infection measurements, and further information for the sensitivity analysis and virtual cohort simulations is also provided (Section S6). A summary of all variables and parameters in the full model (**Eqs. S1-S22**) can be found in **Table S1**.

#### S1. Mathematical model of the immune response to SARS-CoV-2

To model the immune response to SARS-CoV-2 infection, we constructed a system of ordinary and delay differential equations (**Figure 1, Main Text**). The model considers a population of susceptible lung cells ($S$) that become infected ($I$) by SARS-CoV-2 virus ($V$). Infected cells become damaged or dead ($D$) through the virus induced lysis or immune involvement. Upon infection, cells begin secreting type I IFN ($F_U, F_B$)which reduces viral infection and results in cells resistant to virus infection ($R$).

Alveolar macrophages ($M_{\Phi R}$) are activated by infected or dead cells and become inflammatory macrophages ($M_{\Phi I}$) and begin secreting IL-6 ($L_U, L_B$) and GM-CSF ($G_U, G_B$). Monocytes ($M$) are recruited by the presence of infected cells and stimulated by GM-CSF to differentiate into inflammatory macrophages. Neutrophils ($N$) are recruited by G-CSF ($C_U, C_B$) and contribute to bystander death of epithelial cells through the release of reactive oxygen species (ROS). CD8$^+$ T cells ($T$) are recruited after a delay from initial infection and induce apoptosis in infected cells. Bound and unbound concentrations are modelled explicitly. Equations for these interactions are given below:

$$\frac{dV}{dt} = pI - \delta_{V,M\Phi} M_{\Phi I} V - \delta_{V,N} NV - d_V V, \quad S1$$

$$\frac{dS}{dt} = \lambda_S \left(1 - \frac{S+I+R+D}{S_{max}}\right) S - \beta SV - \frac{\rho \delta_N S N^{h_N}}{N^{h_N} + {IC_{50,N}}^{h_N}}, \quad S2$$

$$\frac{dI}{dt} = \frac{\beta \epsilon_{F,I}}{F_B + \epsilon_{F,I}} S(t-\tau_I) V(t-\tau_I) - d_I I - \frac{\delta_N I N^{h_N}}{N^{h_N} + {IC_{50,N}}^{h_N}} - \delta_{I,M\Phi} M_{\Phi I} I - \delta_{I,T} T I, \quad S3$$

$$\frac{dR}{dt} = \lambda_S \left(1 - \frac{S+I+R+D}{S_{max}}\right) R + \frac{\beta F_B}{F_B + \epsilon_{F,I}} S(t-\tau_I) V(t-\tau_I) - \frac{\rho \delta_N R N^{h_N}}{N^{h_N} + {IC_{50,N}}^{h_N}}, \quad S4$$

$$\frac{dD}{dt} = d_I I + \frac{\delta_N (\rho S + \rho R + I) N^{h_N}}{N^{h_N} + {IC_{50,N}}^{h_N}} + \delta_{I,M\Phi}(M_{\Phi I}) I + \delta_{I,T} T I - d_D D + (\delta_{M\Phi,\mathrm{D}} - \delta_{D,M\Phi})(M_{\Phi R} + M_{\Phi I}) D, \quad S5$$

$$\frac{dM_{\Phi R}}{dt} = -a_{I,M\Phi} M_{\Phi R}(I+D) - \delta_{M\Phi,\mathrm{D}} D M_{\Phi R} + \left(1 - \frac{M_{\Phi R}}{M_{\Phi max}}\right) \frac{\lambda_{M_\Phi} M_{\Phi I}}{V + \epsilon_{V,M\Phi}} - d_{M\Phi_R} M_{\Phi R}, \quad S6$$

$$\frac{dM_{\Phi I}}{dt} = a_{I,M\Phi} M_{\Phi R}(I+D) + \frac{p_{M_{\Phi I},G} {G_B}^{h_{M,M\Phi}} M}{{G_B}^{h_{M,M\Phi}} + {\epsilon_{G,M_{\Phi I}}}^{h_{M,M\Phi}}} + \frac{p_{M_{\Phi I},L} L_B M}{L_B + \epsilon_{L,M_\Phi}} - d_{M_{\Phi I}} M_{\Phi I} - \delta_{M\Phi,\mathrm{D}} D M_{\Phi I} - \left(1 - \frac{M_{\Phi R}}{M_{\Phi max}}\right) \frac{\lambda_{M_\Phi} M_{\Phi I}}{V + \epsilon_{V,M\Phi}}, \quad S7$$

$$\frac{dM}{dt} = \left(M_{prod}^* + \left(\psi_M^{max} - M_{prod}^*\right) \frac{{G_B}^{h_M}}{{G_B}^{h_M} + {\epsilon_{G,M}}^{h_M}}\right) M_R + \frac{p_{M,I} I M}{I + \epsilon_{I,M}} - \frac{p_{M_{\Phi I},G} {G_B}^{h_{M,M\Phi}} M}{{G_B}^{h_{M,M\Phi}} + {\epsilon_{G,M_{\Phi I}}}^{h_{M,M\Phi}}} - \frac{p_{M_{\Phi I},L} L_B M}{L_B + \epsilon_{L,M_\Phi}} - d_M M, \quad S8$$

$$\frac{dN}{dt} = \left(N_{prod}^* + \left(\psi_N^{max} - N_{prod}^*\right) \frac{C_{BF} - C_{BF}^*}{C_{BF} - C_{BF}^* + \epsilon_{C,N}}\right) N_R + \frac{p_{N,L} L_B}{L_B + \epsilon_{L,N}} - d_N N, \quad S9$$

$$\frac{dT}{dt} = \frac{p_{T,I} I(t - \tau_T) \epsilon_{L,T}}{L_B + \epsilon_{L,T}} + \frac{p_{T,F} F_B T}{F_B + \epsilon_{F,T}} - d_T T, \quad S10$$

$$\frac{dL_U}{dt} = \frac{p_{L,I} I}{I + \eta_{L,I}} + \frac{p_{L,M_{\Phi I}} M_{\Phi I}}{M_{\Phi I} + \eta_{L,M_{\Phi \mathrm{I}}}} + \frac{p_{L,M} M}{M + \eta_{L,M}} - k_{lin_L} L_U - k_{B_L}\big((M + N + T) A_L - L_B\big) L_U + k_{U_L} L_B, \quad S11$$

$$\frac{dL_B}{dt} = -k_{int_L} L_B + k_{B_L}\big((M + N + T) A_L - L_B\big) L_U - k_{U_L} L_B, \quad S12$$

$$\frac{dG_U}{dt} = \frac{p_{G,M_{\Phi I}} M_{\Phi I}}{M_{\Phi I} + \eta_{G,M\Phi}} + \frac{p_{G,M} M}{M + \eta_{G,M}} - k_{lin_G} G_U - k_{B_G}(M A_G - G_B) G_U + k_{U_G} G_B, \quad S13$$

$$\frac{dG_B}{dt} = -k_{int_G} G_B + k_{B_G}(M A_G - G_B) G_U - k_{U_G} G_B, \quad S14$$

$$\frac{dC_U}{dt} = \frac{p_{C,M} M}{M + \eta_{C,M}} - k_{lin_C} C_U - k_{B_C}(N A_C - C_B)(C_U)^{POW} + k_{U_C} C_B, \quad S15$$

$$\frac{dC_B}{dt} = -k_{int_C} C_B + k_{B_C}(N A_C - C_B)(C_U)^{POW} - k_{U_C} C_B, \quad S16$$

$$\frac{dF_U}{dt} = \frac{p_{F,I} I}{I + \eta_{F,I}} + \frac{p_{F,M_{\Phi I}} M_{\Phi I}}{M_{\Phi I} + \eta_{F,M_{\Phi I}}} + \frac{p_{F,M} M}{M + \eta_{F,M}} - k_{lin_F} F_U - k_{B_F}\big((T + I) A_F - F_B\big) F_U + k_{U_F} F_B, \quad S17$$

$$\frac{dF_B}{dt} = -k_{int_F} F_B + k_{B_F}\big((T + I) A_F - F_B\big) F_U - k_{U_F} F_B, \quad \mathrm{S18}$$

where

$$A_L = \frac{MM_L}{6.02214 \times 10^{23}} \left(K_{L,N} + K_{L,T} + K_{L,M}\right) \cdot \left(\frac{10^{-3}}{5000}\right), \quad S19$$

$$A_G = \frac{MM_G}{6.02214 \times 10^{23}} K_{G,M} \cdot \left(\frac{10^{-3}}{5000}\right), \quad S20$$

$$A_C = \hat{p} \frac{MM_C}{6.02214 \times 10^{23}} K_{C,N} \cdot \left(\frac{10^{1}}{5000}\right), \quad S21$$

$$A_F = \frac{MM_F}{6.02214 \times 10^{23}} \left(K_{F,T} + K_{F,I}\right) \cdot \left(\frac{10^{-3}}{5000}\right). \quad S22$$

#### S2. Model parameter values

A summary of each parameter value in **Eqs. S1-S22** is provided in **Table S1** with references. If the value was obtained through fitting, a reference to the relevant figure is provided. A summary of the variables in the model is given at the end of **Table S1**.

***Table S1 Parameter values used in the Main Text.*** *Parameters have been grouped into: (a-e) cell related, (f-k) cytokine related parameters (l) and initial conditions. Relevant references are given estimated parameters. Parameters obtained through fitting to data in the literature have the appropriate figure noted in the Info column. Parameters estimated from homeostasis calculation are denoted by H or qualitatively estimated by E. Parameters whose value was taken from another parameters estimated has that parameter noted. Viral load is reported as virion copies and cells have been noted in* $10^9$*cells. Time* $t$ *is in days. The final sub-table (m) is a list of the variables in the model.*

*a) Viral kinetic parameters*

| Param | Units | Description | Value | Ref | Info |
|---|---|---|---|---|---|
| $p$ | 1/day ×cop/ml /$10^9$cells | Lytic viral production rate | 741 | 1 | Fig 2 |
| $\lambda_S$ | 1/day | Proliferation of epithelial cells | 0.74 | 2 | Fig S2 |
| $S_{max}$ | $10^9$cells | Epithelial cells carrying capacity | $S_0$ | 3 | |
| $\lambda_{M\Phi}$ | cop/ml/day | Production of alveolar macrophages | 5943 | 4 | Fig S3 |
| $M_{\Phi max}$ | $10^9$cells/ml | Alveolar macrophage carrying capacity | $M_{\Phi R,0}$ | | Fig S3 |
| $\beta$ | 1/day× 1/log(cop/ml) | SARS-CoV-2 virus infection rate | 0.29 | 1 | Fig 2 |
| $\tau_I$ | day | Eclipse time | 0.17 | 5 | |
| $d_I$ | 1/day | Death rate of infected cells | 0.014 | | Fig 2 |
| $\tau_T$ | 1/day | Delay in CD8$^+$ T cell arrival | 4.5 | 6 | |

*b) Cell production, recruitment, and activation rates*

| Param | Units | Description | Value | Ref | Info |
|---|---|---|---|---|---|
| $p_{M\Phi I,G}$ | 1/day | Monocyte to macrophage differentiation by GM-CSF | 1.7 | 7 | |
| $p_{M\Phi I,L}$ | 1/day | Monocyte to macrophage differentiation by IL-6 | 1.7 | 7 | |
| $a_{I,M\Phi}$ | ml/($10^9$cells)× (1/day) | Activation of macs by infected and dead cells | $1.1 \times 10^3$ | 8,9 | |

| | | | | | |
|---|---|---|---|---|---|
| $p_{M,I}$ | 1/day | Monocyte recruitment rate by infected cells | 0.22 | [10] | |
| $p_{T,F}$ | 1/day | $CD8^+$ T cell production rate by IFN | 4 | [11] | |
| $p_{N,L}$ | 1/day | Neutrophils recruitment rate by IL-6 | 0.21 | | H |
| $p_{T,L}$ | 1/day | $CD8^+$ T cells recruitment rate by IL-6 | 4 | [11] | |
| $p_{T,I}$ | 1/day | $CD8^+$ T cell proliferation rate | 1 | [12] | |
| $M^*_{prod}$ | 1/day | Homeostasis reservoir release rate | 0.13 | | H |
| $\psi_M^{max}$ | 1/day | Maximal reservoir release rate | 11.55 | [13] | |
| $N^*_{prod}$ | 1/day | Homeostasis reservoir release rate | 0.21 | | H |
| $\psi_N^{max}$ | 1/day | Maximal reservoir release rate | 4.13 | [14] | |
| $C^*_{BF}$ | — | Homeostasis neutrophil receptor bound fraction | $1.6 \times 10^{-5}$ | [14] | |

*c) Cell-related half-effect ($\epsilon$), IC50 ($IC_{50}$), and Hill coefficient ($h$) parameters*

| **Param** | **Units** | **Description** | **Value** | **Ref** | **Info** |
|---|---|---|---|---|---|
| $\epsilon_{F,I}$ | pg/ml | IFN inhibition of viral production | $4.7\times 10^{-4}$ | [15] | |
| $\epsilon_{L,M_\Phi}$ | pg/ml | IL-6 monocytes to macrophages | 0.011 | [16] | Fig S1 |
| $\epsilon_{G,M_{\Phi I}}$ | pg/ml | GM-CSF monocyte to macrophages | 0.027 | [17] | Fig S1 |
| $\epsilon_{G,M}$ | pg/ml | GM-CSF recruitment of monocytes | 57.2 | [18] | Fig S1 |
| $\epsilon_{F,T}$ | pg/ml | IFN production of $CD8^+$ T cells | 0.004 | [19] | Fig S1 |
| $\epsilon_{C,N}$ | unitless | G-CSF recruitment of neutrophils | $1.89\times 10^{-4}$ | [14] | |
| $\epsilon_{L,N}$ | pg/ml | IL-6 recruitment of neutrophils | 57.2 | | $\epsilon_{G,M}$ |
| $\epsilon_{I,M}$ | $10^9$cells/ml | Infected cell monocyte recruitment | 0.11 | | E |
| $\epsilon_{L,T}$ | pg/ml | IL-6 production of $CD8^+$ T cells | $3 \times 10^{-4}$ | [21] | Fig S1 |
| $\epsilon_{V,M\Phi}$ | log(cop/ml) | Viral load for mac replenishing | 2.96 | | Fig S3 |
| $\epsilon_{T,I}$ | $10^9$cells/ml | Antigen driven proliferation | $10^{-6}$ | [12] | |
| $h_M$ | — | GM-CSF monocyte recruitment | 1.67 | [18] | Fig S1 |
| $h_{M,M\Phi}$ | — | GM-CSF monocyte to macrophages | 2.03 | [17] | Fig S1 |
| $h_N$ | — | Neutrophil induced damage | 3.02 | [22] | Fig S1 |
| $IC_{50,N}$ | $10^9$cells/ml | Neutrophil induced damage | 0.047 | [22] | Fig S1 |

*d) Cell/virus-induced death rates*

| **Param** | **Units** | **Description** | **Value** | **Ref** | **Info** |
|---|---|---|---|---|---|
| $\delta_{V,M\Phi}$ | ml/($10^9$cells)× 1/day | Rate of viral clearance by macrophages | 768 | [23] | Fig S2 |
| $\delta_{V,N}$ | ml/($10^9$cells)× 1/day | Rate of viral clearance by neutrophils | 2304 | | |
| $\delta_N$ | 1/day | Rate of neutrophil inflicted damage | 1.68 | [24] | Fig S2 |
| $\rho$ | — | Bystander death modulation constant | 0.5 | | E |
| $\delta_{I,M\Phi}$ | ml/($10^9$cells)× (1/day) | Rate macrophages phagocytose infected cells | 121 | [25] | |
| $\delta_{I,T}$ | ml/($10^9$cells)× (1/day) | Rate $CD8^+$ T cells induce apoptosis in infected cells | 238 | [26] | |
| $\delta_{M\Phi,D}$ | ml/($10^9$cells)× (1/day) | Rate macrophages die from phagocytosis | 6.06 | [27,28] | |
| $\delta_{D,M\Phi}$ | ml/($10^9$cells)× (1/day) | Rate macrophages phagocytose dead cells | 8.03 | [27] | Fig S2 |

*e) Cell death and virus decay rates*

| Param | Units | Description | Value | Ref | Info |
|---|---|---|---|---|---|
| $d_V$ | 1/day | Viral decay rate | 1.81 | [29] | Fig 2 |
| $d_D$ | 1/day | Degradation rate of apoptosed cells | 8 | [30] | |
| $d_{M_{\Phi R}}$ | 1/day | Alveolar macrophage death rate | 0 | [31] | |
| $d_{M_{\Phi I}}$ | 1/day | Inflammatory macrophage death rate | 0.3 | [32] | |
| $d_M$ | 1/day | Monocyte death rate | 0.76 | [33] | |
| $d_N$ | 1/day | Neutrophil death rate | 1.28 | [34] | |
| $d_T$ | 1/day | $CD8^+$ T cell death rate | 0.4 | [35] | |

*f) Cytokine production rates*

| Param | Units | Description | Value | Ref | Info |
|---|---|---|---|---|---|
| $p_{L,I}$ | pg/ml/day | IL-6 production by infected cells | 11.89 | [36] | Fig S4 |
| $p_{L,M_{\Phi I}}$ | pg/ml/day | IL-6 production by activated macrophages | 1872 | [37] | Fig S4 |
| $p_{L,M}$ | pg/ml/day | IL-6 production by monocytes | 72.56 | [38] | Fig S1 |
| $p_{G,M_{\Phi I}}$ | pg/ml/day | GM-CSF production by inflammatory macrophages | 2626 | [39] | |
| $p_{C,M}$ | ng/ml/day | G-CSF production by monocytes | 26.26 | | $p_{G,M_{\Phi I}}$ |
| $p_{G,M}$ | pg/ml/day | GM-CSF production by monocytes cells | 3070 | [40] | Fig S3 |
| $p_{F,I}$ | pg/ml/day | IFN production by infected cells | 2.82 | [41] | Fig S3 |
| $p_{F,M_{\Phi I}}$ | pg/ml/day | IFN production by inflammatory macrophages | 1.3 | | E |
| $p_{F,M}$ | pg/ml/day | IFN production by monocytes | 3.56 | [42,43] | Fig S3 |

*g) Cytokine production half-effect (η) and Hill coefficient (h) parameters*

| Param | Units | Description | Value | Ref | Info |
|---|---|---|---|---|---|
| $\eta_{L,I}$ | $10^9$cells/ml | IL-6 production by infected cells | 0.7 | [36] | Fig S4 |
| $\eta_{L,M}$ | $10^9$cells/ml | IL-6 by monocytes | 0.0045 | [38] | Fig S1 |
| $\eta_{L,M_{\Phi I}}$ | $10^9$cells/ml | IL-6 by inflammatory macrophages | $3.6\times 10^{-5}$ | [37] | Fig S4 |
| $\eta_{G,M\Phi}$ | $10^9$cells/ml | GM-CSF by macrophages | $3.6\times 10^{-5}$ | | $\eta_{L,M\Phi}$ |
| $\eta_{G,M}$ | $10^9$cells/ml | GM-CSF by monocytes | 0.15 | | H |
| $\eta_{C,M}$ | $10^9$cells/ml | G-CSF by monocytes | 3.05 | | H |
| $\eta_{F,I}$ | $10^9$cells/ml | IFN by infected cells | 0.011 | [41] | Fig S4 |
| $\eta_{F,M_{\Phi I}}$ | $10^9$cells/ml | IFN by inflammatory macrophages | $1.3\times 10^{-6}$ | | H |
| $\eta_{F,M}$ | $10^9$cells/ml | IFN by monocytes | 0.54 | [42,43] | Fig S3 |

*h) Cytokine linear (renal) clearance and internalization rates*

| Param | Units | Description | Value | Ref | Info |
|---|---|---|---|---|---|
| $k_{lin_L}$ | 1/day | Rate of IL-6 renal clearance | 16.6 | [44] | |
| $k_{lin_G}$ | 1/day | Rate of GM-CSF renal clearance | 11.7 | [45] | |
| $k_{lin_C}$ | 1/day | Rate of G-CSF renal clearance | 0.16 | [14] | |
| $k_{lin_F}$ | 1/day | Rate of IFN renal clearance | 18 | [46] | |
| $k_{int_L}$ | 1/day | Internalization rate of IL-6 | 61.8 | [47] | Fig S2 |
| $k_{int_G}$ | 1/day | Internalization rate of GM-CSF | 73.4 | [48] | |
| $k_{int_C}$ | 1/day | Internalization rate of G-CSF | 462 | [14] | |
| $k_{int_F}$ | 1/day | Internalization rate of IFN | 17 | [49] | |

*i) Cytokine binding/unbinding rates and stoichiometric constant*

| Param | Units | Description | Value | Ref | Info |
|---|---|---|---|---|---|
| $k_{B_L}$ | ml/pg/day | IL-6 binding rate | 0.0018 | [50] | |
| $k_{B_G}$ | ml/pg/day | GM-CSF binding rate | 0.0021 | [48] | |
| $k_{B_C}$ | ml/ng/day | G-CSF binding rate | 2.24 | [14] | |
| $k_{B_F}$ | ml/pg/day | IFN binding rate | 0.011 | [49] | |
| $k_{U_L}$ | 1/day | IL-6 unbinding rate | 22.3 | [50] | |
| $k_{U_G}$ | 1/day | GM-CSF unbinding rate | 522 | [48] | |
| $k_{U_C}$ | 1/day | G-CSF unbinding rate | 184 | [14] | |
| $k_{U_F}$ | 1/day | IFN unbinding rate | 6.07 | [49] | |
| $POW$ | — | Stoichiometric constant (G-CSF) | 1.4608 | [51] | |
| | | Stoichiometric constant (IL-6, GM-CSF, IFN) | 1 | | |
| $\hat{p}$ | — | Stoichiometry relating constant (G-CSF) | 2 | [51] | |
| | | Stoichiometry relating constant (IL-6, GM-CSF, IFN) | 1 | | |

*j) Number of cellular receptors and cytokine molecular weights*

| Param | Units | Description | Value | Ref | Info |
|---|---|---|---|---|---|
| $K_{L,N}$ | sites/cell | No. IL-6 receptors on neutrophils | 720 | [52] | |
| $K_{L,T}$ | sites/cell | No. IL-6 receptors on T cells | 300 | [53] | |
| $K_{L,M}$ | sites/cell | No. of IL-6 receptors on monocytes | 509 | [54] | |
| $K_{G,M}$ | sites/cell | No. GM-CSF receptors on monocyte | 1058 | [55] | |
| $K_{C,N}$ | sites/cell | No. of G-CSF receptors on neutrophil | 600 | [56] | |
| $K_{F,T}$ | sites/cell | No. of IFN receptors on T cells | 1000 | [57] | |
| $K_{F,I}$ | sites/cell | No. of IFN receptors on infected cells | 1300 | [58] | |
| $MM_L$ | g/mol | Molecular weight of IL-6 | 21000 | [59] | |
| $MM_G$ | g/mol | Molecular weight of GM-CSF | 14000 | [60] | |
| $MM_C$ | g/mol | Molecular weight of G-CSF | 19600 | [14] | |
| $MM_F$ | g/mol | Molecular weight of IFN-$\beta$ | 19000 | [61] | |

*k) Initial conditions*

| Param | Units | Description | Value | Ref |
|---|---|---|---|---|
| $V_0$ | $\log_{10}$(copies /ml) | Initial viral load | 12 | [1] |
| $S_0$ | $10^9$ cells/ml | Initial susceptible cells | 0.16 | [3,62] |
| $I_0$ | $10^9$ cells/ml | Initial infected cells | 0 | |
| $R_0$ | $10^9$ cells/ml | Initial resistant cells | 0 | |
| $M_{\Phi R,0}$ | $10^9$ cells/ml | Initial resident macrophages | $2.7 \times 10^{-5}$ | [3] |
| $M_{\Phi I,0}$ | $10^9$ cells/ml | Initial inflammatory macrophages | $2.9 \times 10^{-7}$ | H |
| $M_0$ | $10^9$ cells/ml | Initial monocytes | 0.0004 | [63] |
| $M_R$ | $10^9$ cells/ml | Initial reservoir monocytes | 0.0023 | |
| $N_0$ | $10^9$ cells/ml | Initial neutrophils | 0.0053 | [14] |
| $N_R$ | $10^9$ cells/ml | Initial reservoir neutrophils | 0.0316 | [14] |
| $T_0$ | $10^9$ cells/ml | Initial $CD8^+$ T cells | $1.1 \times 10^{-4}$ | [64,65] |
| $L_{U,0}$ | pg/ml | Initial unbound IL-6 | 1.1 | [66] |
| $L_{B,0}$ | pg/ml | Initial bound IL-6 | $1.4 \times 10^{-6}$ | H |
| $G_{U,0}$ | pg/ml | Initial unbound GM-CSF | 2.43 | [67] |
| $G_{B,0}$ | pg/ml | Initial bound GM-CSF | $1.6 \times 10^{-8}$ | H |

| | | | | |
|---|---|---|---|---|
| $C_{U,0}$ | ng/ml | Initial unbound G-CSF | 0.025 | [14] |
| $C_{B,0}$ | ng/ml | Initial bound G-CSF | $6.5\times 10^{-10}$ | [14] |
| $F_{U,0}$ | pg/ml | Initial unbound IFN | 0.015 | [68] |
| $F_{B,0}$ | pg/ml | Initial bound IFN | $1.1\times 10^{-8}$ | H |

*m) List of variables in* ***Eqs.*** **S1-S22**

| **Variable** | **Units** | **Description** |
|---|---|---|
| $V$ | cop/ml | Viral load |
| $S$ | $10^9$ cells/ml | Susceptible cells |
| $I$ | $10^9$ cells/ml | Infected cells |
| $R$ | $10^9$ cells/ml | Resistant cells |
| $M_{\Phi R}$ | $10^9$ cells/ml | Alveolar (resident) macrophages |
| $M_{\Phi I}$ | $10^9$ cells/ml | Inflammatory macrophages |
| $M$ | $10^9$ cells/ml | Monocytes |
| $M_R$ | $10^9$ cells/ml | Bone marrow reservoir monocytes |
| $N$ | $10^9$ cells/ml | Neutrophils |
| $N_R$ | $10^9$ cells/ml | Bone marrow reservoir neutrophils |
| $T$ | $10^9$ cells/ml | $CD8^+$ T cells |
| $L_U$ | pg/ml | Unbound IL-6 |
| $L_B$ | pg/ml | Bound IL-6 |
| $G_U$ | pg/ml | Unbound GM-CSF |
| $G_B$ | pg/ml | Bound GM-CSF |
| $C_U$ | ng/ml | Unbound G-CSF |
| $C_B$ | ng/ml | Bound G-CSF |
| $C_{BF}$ | unitless | Neutrophil G-CSF receptor bound fraction |
| $F_U$ | pg/ml | Unbound IFN |
| $F_B$ | pg/ml | Bound IFN |

### S3. Parameters taken from literature

#### *S3.1. Initial cytokine concentrations*

The basal concentrations of unbound cytokines were taken from values in the literature. The plasma concentration of colony stimulating factor GM-CSF in healthy adults was measured by Lee et al.[67] to be $2.43 \pm 0.42$ pg/ml (i.e., $G_{U,0} = 2.43$ pg/ml). We fixed the initial unbound G-CSF cytokine concentration to $C_{U,0} = 0.025$ ng/ml [14]. The concentration of unbound IFN type 1 was set to be $F_{U,0} = 0.015$ pg/ml based on the average value of IFN-α in humans[69,70]. The median plasma IL-6 concentration was estimated to be $L_{U,0} = 1.1$ pg/ml in blood samples from healthy adults[71].

#### *S3.2. Initial cell populations*

The average total number of type I and type II alveolar epithelial cells and endothelial cells in the lung was estimated by Crapo et al.[3] to be $136 \times 10^9$ cells from eight people (6 males, 2 females) aged 19-40,. At functional residual capacity, pulmonary total tissue volume was reported by Armstrong et al.[62] to be $843 \pm 110$ ml. Together, this gave an initial target cell concentration of $S_0 = 0.16 \times 10^9$ cells/ml. Similarly, Crapo et al.[3] found the average number of alveolar macrophages to be $23 \pm 7 \times 10^6$ cells, thus $M_{\Phi R,0} = 2.73 \times 10^5$ cells/ml. Monocytes account for 1% to 10% of circulating white blood cells, which equates to 200 to 600 monocytes per microliter of blood[63] (with a blood volume of 5 litres[72]). Therefore, we assumed that at homeostasis $M_0 = 4 \times 10^5$ cells/ml. For the total number of neutrophils in the blood, we used the previous estimate of Craig et al.[51] $N_0 = 5.26 \times 10^6$ cells/ml that was calculated from whole blood and marginated neutrophils. Lastly, the number of $CD8^+$ T cells in the lung tissue was, on average, 20% of the number of $CD8^+$ T cells in the blood[65]. Using flow cytometry, Uppal et al.[64] determined there were $552$ cells/μl, on average, in the blood. To account for the number of naïve $CD8^+$ T cells infiltrating the lungs[73], the initial number of $CD8^+$ T cells was estimated from the proportion of T cells in the tissue, i.e. $T_0 = 1.1 \times 10^5$ cells/ml.

#### *S3.3. Cytokine molecular weights and receptors per cell*

Cytokine binding and unbinding kinetics were modelled using the molecular weight of each cytokine and the number of corresponding receptors on the binding cell. The molecular weight for IL-6 is $MM_L = 21{,}000$ g/mol[59]; for GM-CSF is $MM_G = 14{,}000$ g/mol[60]; for G-CSF is $MM_C = 19{,}600$ g/mol[51]; and for IFN-β is $MM_F = 19{,}000$ g/mol[61]. The number of high-affinity receptors for GM-CSF on the surface of blood monocytes[55] is $K_{G,M} = 1{,}058$ sites/cell. Mature human neutrophils express ~200-1,000 G-CSF receptors per cell[56], thus we fixed $K_{C,N} = 600$ sites/cell. Most cells have 1,000-2,000 type I IFN receptors (IFNAR) receptors[57]. Assuming $CD8^+$ T cells are at the lower end of this interval gives $K_{F,T} = 1{,}000$ sites/cell. The number of IFNAR sites on HEC1B human (uterus/endometrium epithelial) cells is on average $K_{F,I} = 1{,}300$ sites/cell[58]. The number of IL-6 receptors on $CD8^+$ T cells is $K_{L,T} = 300$ sites/cell based on measurements from Taga et al.[53]. We fixed the number of IL-6 receptors on neutrophils and monocytes to $K_{L,N} = 720$ sites/cell and $K_{L,M} = 509$ sites/cell respectively, based on the range in IL-6 receptors expressed on myeloma hematopoietic cells[52] and IL-6 receptors on mouse myelomonocytic leukemic M1 cells[54].

#### *S3.4. Binding/unbinding rates of cytokines*

Based on measurements by Tenhumberg et al.[50], the binding and unbinding rate of IL-6 was set to be $k_{B_L} = 0.0018$ pg/ml/day and $k_{U_L} = 22.29$/day, respectively. The unbinding kinetics for G-CSF were previously estimated by Craig et al.[51] to be $k_{U_C} = 184.87$/day. Lastly, the unbinding and binding rates for GM-CSF were taken from previous modelling work[48] and set to $k_{U_G} = 522.72$/day and $k_{B_G} = 0.0021$ per pg/ml/day respecitvely. Mager and Jusko[49] estimated the binding rate of IFN-β using a PKPD model to be $k_{B_F} = 0.0107$ per pg/ml/day. Lastly, the binding rate for GM-CSF was fixed as $k_{B_G} = 0.0021$ per pg/ml/day[48].

#### *S3.5. Clearance and internalization rate of cytokines*

Most of the clearance and internalization rates of cytokines in **Eqs. S1-S22** were obtained by assuming exponential clearance and using the half-life formula:

$$k_{lin} = \frac{\ln(2)}{t_{1/2}}, \qquad S23$$

where $t_{1/2}$ is the cytokine half-life and $k_{lin}$ the clearance rate. IL-6 has a short half-life of approximately 1 hour[44] giving a clearance rate of $k_{lin_L} = 16.6$/day. The half-life of GM-CSF ranges between 50-85 minutes[45]. We took the upper value giving $k_{lin_G} = 11.74$/day. The linear clearance rate of G-CSF was previously estimated by Craig et al.[51] as $k_{lin_C} = 0.16$/day. Half-lives for IFN-β range from 1-2 hours[46]. Using the lower bound, we fixed the clearance rate as $k_{lin_F} = 16.6$/day, giving a half-life of roughly 55 minutes. The internalization rates of GM-CSF and G-CSF were taken from previous pharmacokinetic modelling[48,51] and fixed to $k_{int_G} = 73.4$/day and $k_{int_C} = 462$/day. Similarly, the internalization rate of IFN was fixed as $k_{int_F} = 16.97$/day based on previous modelling of the receptor-mediated dynamics of IFN-β[46,74]. The internalization rate of IL-6, $k_{lin_L}$, was estimated by data fitting (Section S4.2.2).

#### *S3.6. Neutrophil and monocyte reservoir dynamics*

Craig et al.[51] previously developed a physiological model of the production dynamics of neutrophils through G-CSF regulation that accounts for the concentration of freely circulating cytokine and cytokine bound to mature neutrophils. The parameters in our model that relate to the number of neutrophils in the bone marrow reservoir, release rate, and the dynamics of G-CSF on neutrophils (**Eq. 5** & **Eq. S9**) were taken from their work, i.e. $N_R = 3.16 \times 10^7$ cells/ml, $C_{BF}^* = 1.58 \times 10^{-5}$ (unitless), $\epsilon_{C,N} = 1.8924 \times 10^{-4}$ (unitless), and $\psi_N^{max} = 4.13/\text{day}$. Similarly, the bone marrow monocyte reservoir dynamics were estimated based on previous modelling work by Cassidy et al.[13] to be $M_R = 2.27 \times 10^6$ cells/ml and $\psi_M^{max} = 11.55/\text{day}$.

#### *S3.7. Monocyte, macrophage differentiation and activation rates*

Previous mathematical modelling studies were used to estimate the monocyte and macrophage differentiation and activation rates. We assumed that the recruitment rate of monocytes would be equal to the recruitment rate of new macrophages by infected cells[10], giving $p_{M,I} = 0.22/\text{day}$. We approximated the activation rate of resident macrophages to inflammatory macrophages from the rate of dendritic cell activation[8,9], giving $a_{I,M\Phi} = 1.1 \times 10^3$ per $10^9$ cells/ml/day. Lastly, the GM-CSF and IL-6 stimulated differentiation rate of monocytes to macrophages was $p_{M_{\Phi I},G} = 1.7$/day based on the estimates that it can take 12-14 hours for monocytes to migrate from the bone marrow to the site of inflammation and subsequently differentiate into progenitor cells[7].

#### *S3.8. CD8$^+$ T cell recruitment and expansion rate*

The maximal time for CD8$^+$ T cell division is between 4-6 hours[11], giving a production rate of $p_{T,F} =$ 4/day. The dynamics of CD8$^+$ T cells in response to infected cells were modelled similarly to previous work by de Pillis et al.[75] and Baral et al.[12], and we set the activation rate and half-effect parameter of infected cells from these studies, i.e. $p_{T,I} = 9 \times 10^{-3}$/day and $\epsilon_{T,I} = 10^3$ cells/ml. Lastly, $\tau_T = 4.5$ days based on the delay in infected cell recruitment of CD8$^+$ T cells[6].

#### *S3.9. Cell death rates*

Neutrophils are known to have a short half-life in circulation[34] of $d_N = 1.28$/day. Kim et al.[35] estimated that primed CD8$^+$ T cells have a death rate of $d_T = 0.4$/day. Monocytes transiting from the bone marrow to the blood have a circulating half-life of 22 hours[33], using the half-life formula (**Eq. S23**) this gives $d_M = 0.756/\text{day}$. The time from initiation of cell apoptosis to completion can occur as quickly as 2-3 hours[30] giving a dead cell decay rate of $d_D = 8$/day. At homeostasis, mature macrophages are a quiescent population with a half-life between 4-6 weeks[31], or sometimes greater than 80 days[76]. Thus, we assumed that the death of resident alveolar macrophages is negligible ($d_{M\Phi R} = 0$/day) given the time frame of acute SARS-CoV-2 infections considered in this study (3 weeks). Inflammatory macrophages were assumed to have a death rate of $d_{M_{\Phi I}} = 0.3$/day, which was estimated through inflammatory macrophages in response to oncolytic virotherapy[32]. Macrophages also undergo apoptosis from phagocytosing too much material (exhaustion). We assumed it takes ~20 dead cells to be phagocytosed to induce macrophage death[27,28] and set $\delta_{M\Phi,D} = 6.06$ per $10^9$ cells/day. The rate of CD8$^+$ T cell-induced apoptosis of infected cells was previously estimated by Lee et al.[26] giving $\delta_{I,T} = 238$ per $10^9$ cells/day. The phagocytosis rate of infected cells by macrophages $\delta_{I,M\Phi} = 121.195$ per $10^9$ cells/day was estimated from neutrophil phagocytosis[25].

### S4. Parameters estimated by data fitting

#### S4.1. Pharmacodynamics of stimulatory and inhibitory effects for cells and cytokines

We used standard pharmacodynamic relationships to model the various immunological effects of cells and cytokines. Here, the half-maximal response (generally expressed as an $EC_{50}$ or $IC_{50}$) of the cytokine or cell population is the concentration at which half of the maximal (stimulatory of inhibitory) effect is achieved[77]. Effect curves, $E$, (stimulatory or inhibitory respectively) are given by[78]

$$E = E_0 + E_{max} \frac{W^h}{W^h + EC_{50}^h}, \qquad S24$$

$$E = E_0 + E_{max} \left(1 - \frac{W^h}{W^h + {IC_{50}}^h}\right), \qquad S25$$

where $E$ denotes the measured response (e.g., cell viability), $W$ is the concentration of cytokine or cells under consideration, $E_0$ is the basal effect (the response when the dose of the compound is zero), $E_{max}$ is the maximum effect (stimulatory of inhibitory) of the compound, $h > 0$ is the Hill coefficient that measures the sensitivity of the response to the dose range of the compound (i.e. the slope of the dose-response curve). **Eqs. S24-S25** are also known as Emax and Imax functions[77], see also **Eq. 1 Main Text**.

##### S4.1.1. *Type I IFN inhibition of viral infection and replication*

Sheahan et al.[15] measured the mean inhibition of MERS-CoV replication of IFN-β and found the $EC_{50}$ to be 175 IU/ml. The specific activity of recombinant human IFN-β is approximately $2.8 \times 10^8$ IU/mg[15]. We converted the $EC_{50}$ of 175 IU/ml to pg/ml to give the half-effect of IFN-β on viral production and infection capacity as $\epsilon_{F,I} = 625$ pg/ml[15]. Since this is a measurement of unbound IFN, we scaled this by the initial proportion of bound to unbound IFN ($F_{B,0}/F_{U,0}$) to get $\epsilon_{F,I} = 4.65 \times 10^{-4}$ pg/ml.

##### S4.1.2. *Neutrophil-induced damage of alveolar epithelial cells*

To estimate the rate of neutrophil induced damage due to the release of reactive oxygen species (ROS) (**Eqs. S2-S4**), we used cell viability measurements of rat alveolar epithelial cells (RLE) after incubation for 2 hours with hydrogen peroxide ($H_2O_2$) at varying concentrations[22]. Fitting **Eq. S25** to this data, we obtained $IC_{50} = 197.63$µM and $h = 3.02$ (**Figure S1**A). To convert this $IC_{50}$ from a concentration of $H_2O_2$ to the concentration of neutrophils (as in **Eqs. S2-S4**), we used the approximate amount of $H_2O_2$ produced by a single neutrophil in response to stimulation by *N*-Formylmethionyl-leucyl-phenylalanine and phorbol myristate acetate[79]. Taking the average response from these stimuli and converting to units µM/cell, neutrophils produce 0.0042µM/cell of $H_2O_2$. As the maximum production of $H_2O_2$ by neutrophils is achieved relatively fast (15-30 minutes after stimulation[79]) we estimated the equivalent $IC_{50}$ as a neutrophil concentration to be $4.71 \times 10^4$ cells (i.e. $197.62/0.0042$). As this represents a concentration of stimulated neutrophils we then increased this by the number of neutrophils at homeostasis[80] to give the half-effect concentration for neutrophil bystander damage of $IC_{50,N} = 4.71 \times 10^7$cells/ml. This estimate was in line with estimates obtained from similar experiments conducted by Weiss et al[81] and Snyers et al.[52].

##### S4.1.3. *Effect of GM-CSF on monocyte production and differentiation*

GM-CSF can act in a paracrine fashion to recruit circulating monocytes, enhance their functions in host defense[82,83] and influence their differentiation into monocytic or granulocytic lineages[17]. Reducing **Eqs. S8** to consider only the effect of production and differentiation gives

$$\frac{dM}{dt} = \frac{p_{M,G} {G_B}^{h_M}}{{G_B}^{h_M} + {\epsilon_{G,M}}^{h_M}} - \frac{p_{M_{\Phi I},G} {G_B}^{h_{M,M\Phi}} M}{{G_B}^{h_{M,M\Phi}} + {\epsilon_{G,M_{\Phi I}}}^{h_{M,M\Phi}}}. \quad S26$$

We modelled the effect of GM-CSF on monocyte production by estimating its potency from the dose-response of cultured blood monoculture cells with various concentrations of murine recombinant GM-CSF for 21 days[18]. Fitting **Eq. S24** gave $EC_{50} = 85.8$ IU/ml and $h = 1.67$ (**Figure S1**B). Using the specific activity of recombinant GM-CSF (i.e. $15 \times 10^5$ IU/µg [18]), this becomes $\epsilon_{G,M} = 57.2$ pg/ml. GM-CSF also promotes myeloid differentiation of cultured bone marrow cells into granulocytic and monocytic lineages towards terminal differentiation into monocytes, macrophages, and dendritic cells[17]. Sun et al.[17] investigated whether the dose of GM-CSF regulates the development of myeloid cells and measured the monocytic myeloid cell count as a function of GM-CSF concentrations. Fitting **Eq. S24** to this data, we obtained $EC_{50} = 2.66$ ng/ml and $h = 2.03$ (**Figure S1**C). Converting this to the units of GM-CSF in our model gives $2.7 \times 10^3$ pg/ml as the unbound GM-CSF half-effect concentration, which scaled by $10^{-5}$ (the order of initial bound GM-CSF to initial monocytes $G_{B,0}/M_0$) gives a bound GM-CSF half effect concentration of $\epsilon_{G,M\Phi} = 0.027$ pg/ml.

##### S4.1.1. *IL-6 production by monocytes and effect on monocyte differentiation*

Peripheral blood monocytes can be induced to secrete an array of cytokines, including IL-6, by stimuli such as lipopolysaccharide (LPS)[38]. The production of unbound IL-6 was modelled as a function of the monocyte concentration (**Eq. S11**):

$$\frac{dL_U}{dt} = \frac{p_{L,M} M}{M + \eta_{L,M}}. \quad S27$$

Alderson et al.[38] measured the concentration of IL-6 (IU/ml) produced by monocytes stimulated with 10µl of LPS over 24 hours in 1ml of culture medium (**Figure S1**D). We fit **Eq. S27** to this data by assuming the number of monocytes was fixed to $M = 2 \times 10^5$ cells and that there was no monocyte proliferation over the course of the experiment. This gave an estimate of $p_{L,M} = 7.26 \times 10^4$ pg/ml/day (converted using 4.5 pg/ml as the concentration required for half-maximal stimulation of B9 proliferation by IL-6[84]) and $\eta_{L,M} = 9 \times 10^4$ cells. Scaling this half-effect by the initial concentration of monocytes gives $\eta_{L,M} = 4.98 \times 10^7$ cells/ml. We then scaled the production rate of IL-6 by $10^3$ as the maximum IL-6 concentration achieved *in vivo* during SARS-CoV-2 infection was $10^3$ less than that in the two *in vitro* experiments. We confirmed this production rate using the experiments of Morris et al.[85], which measure the production of IL-6 by blood mononuclear cells by co-culturing with airway smooth muscle cells (ASM) cells and LPS stimulation.

Production of macrophages by monocytes from stimulation with IL-6 was modelled as

$$\frac{dM}{dt} = -\frac{p_{M_{\Phi I},L} L_B M}{L_B + \epsilon_{L,M_\Phi}}, \quad S28$$

$$\frac{dM_{\Phi I}}{dt} = \frac{p_{M_{\Phi I},L} L_B M}{L_B + \epsilon_{L,M_\Phi}}, \quad S29$$

where $p_{M_{\Phi I},I}$ is the production rate and $\epsilon_{L,M\Phi}$ is the half effect from the bound IL-6 (**Eqs. S7-S8**). To estimate the production of macrophages based on the concentration of IL-6, we used measurements for the production of macrophages by fibroblasts. Fibroblasts release IL-6, which then up-regulates the expression of functional M-CSF receptors on monocytes[16] and allows monocytes to consume autocrine M-CSF and thus switch differentiation to macrophages rather than DCs. Chomarat et al.[16] cultured monocytes in GM-CSF and IL-4 with graded numbers of normal skin fibroblasts. At day 5, cells were analyzed for macrophage

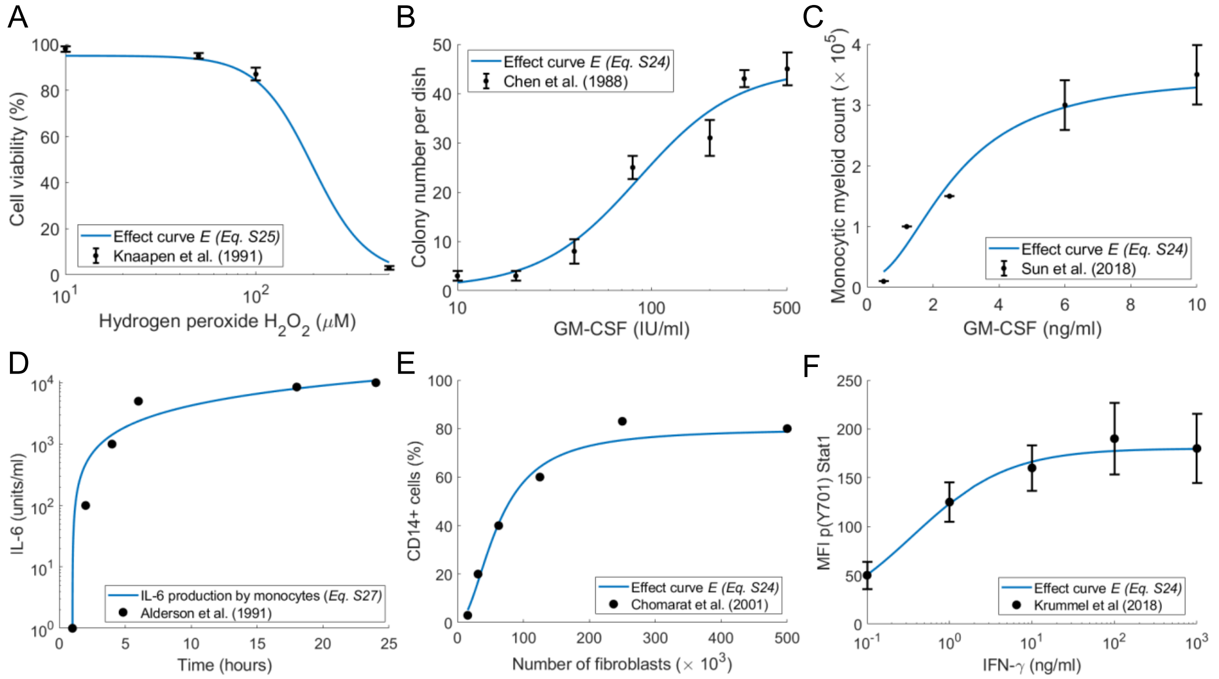


***Figure S1. Effects of neutrophils on lung epithelial cells, GM-CSF on monocyte production and differentiation, the relationships between monocytes and CD4+ T cells with IL-6, and the influence of IFN on T cell expansion.*** ***A)*** *Using the measurements by Knaapen et al.*[22]*, the inhibitory effect curve E (**Eq. S25**) was fit to the cell viability of RLE cells under various concentrations of* $H_2O_2$*.* ***B)*** *The stimulatory effect curve E (**Eq. S24**) was fit to the dose response measurements of blood monoculture cells (*$3 \times 10^3$*cells/dish) with various concentrations of murine recombinant GM-CSF (IU/ml)*[18]*.* ***C)*** *The stimulatory effect curve E (**Eq. S24**) was fit to measurements for the monocytic myeloid cell count as a function of GM-CSF.*[17] ***D) Eq. S27*** *fit to time course data of IL-6 production from monocytes*[38]*.* ***E)*** *IL-6 stimulation of monocyte differentiation to macrophages modelled by the inhibitory effect curve E (**Eq. S24**) fit to the percentage of CD14+ cells (macrophages) as a function of the number of fibroblasts measured by Chomarat et al.*[16]*.* ***F)*** *Stimulatory effect curve E (**Eq. S24**) for IFN-γ stimulation on CD8+ T cells fit to measurements of the signalling in CD8+T cells for varying doses of IFN-γ*[19]*. Data (black) is plotted as either circles (**D & E**) or mean and standard deviation error bars (**A-C&F**); solid blue line: corresponding fit.*

markers CD1a and CD14. Fitting **Eq. S24** to these results gave an $EC_{50} = 61.6$ cells and $h = 1.96$ (**Figure S1**E). Using then concentration of IL-6 produced by 250,000 fibroblasts (4.5pg/ml) and assuming that there is a linear relationship between the number of fibroblasts and the concentration of IL-6, we converted this to an unbound IL-6 concentration, i.e. $EC_{50} = 1.1 \times 10^3$ pg/ml. Scaling this by $10^{-5}$ (the order of initial bound to unbound IL-6 in the model) gives the bound IL-6 concentration $\epsilon_{L,M\Phi} = 0.011$ pg/ml.

##### S4.1.1. *Effect of IFN on CD8+ T cells*

To estimate the half-effect IFN concentration for $CD8^+$ T cell regulation, $\epsilon_{F,T}$, we used dose-response measurements for $CD8^+$ T regulation by IFN-γ. IFN-γ is known to regulate $CD8^+$ T cell differentiation through co-stimulation of the signal transducer and activator of transcription 1 (STAT1) pathway[19]. Krummel et al.[19] measured the effect on IFN-γ signaling in $CD8^+$ T cells by analysing the p(Y701) STAT1 as a function of increasing concentrations. Fitting **Eq. S24** to this data for $EC_{50}$ and $h$ resulting in a $h$ value of approximately 1, so we fixed $h = 1$ to improve identifitability and fit $EC_{50}$ which gave $EC_{50} = 0.4$ ng/ml. Assuming the half-effect concentration for IFN-γ regulation of the STAT1 pathway can be used to estimate the half-effect concentration of IFN gives $\epsilon_{F,T} = 0.004$ pg/ml (scaled by $10^{-5}$ to obtain the unbound concentration; **Figure S1**F).

S4.1.1. ***Effect of IL-6 on CD4*** $^{+}$ ***T cell expansion***

IL-6 stimulates IL-2 production and the proliferation of $CD8^{+}$ and $CD4^{+}$ T cells[21]. Holsti and Raulet[21] measured the counts per minute (CPM) of $CD4^{+}$ cell proliferation from IL-6 and IL-1 induction. Converting their data from sample dilution to µg/ml and fitting **Eq. S24** gave $h = 2$ and $EC_{50} = 2.26 \times 10^{-4}$ (reciprocal dilution of IL-6) (**Figure S2**A). In Holsti and Raulet, the medium contained 2.1 µg/ml of IL-6. Converting the dilution to a concentration in our units and scaling this by $10^{-5}$ (the order of initial bound IL-6 to unbound IL-6) gives $\epsilon_{L,T} = 4.7 \times 10^{-3}$ pg/ml.

#### S4.2. Estimating parameters from temporal data

S4.2.1. ***Proliferation rate of epithelial cells***

We used measurements of A549 cell proliferation[2] to fit an exponential growth curve and determined the proliferation rate of epithelial cells to be $\lambda_S = 0.744$/day (**Figure S2**B).

S4.2.2. ***IL-6 internalization rate***

Bound IL-6 is internalized at a rate $k_{int_L} L_B$ (**Eq. S12**), which gives the fraction of internalized IL-6

$$f(t) = 1 - e^{-k_{int_L} t}. \qquad S30$$

Fitting $k_{int_L}$ to experiments measuring the fraction of IL-6 internalized by hepatocytes over 30 minutes[47] gave $k_{int_L} = 61.8$/day (**Figure S2**C).

S4.2.3. ***Neutrophil-induced cell death rate***

To estimate the rate of epithelial cell death induced by the release of $H_2O_2$ by neutrophils ($\delta_N$), we used measurements of the total alveolar macrophages apoptosis after $H_2O_2$ exposure from 0-12 hours[24] and fit an exponential decay rate, which gave $\delta_N = 1.68$/day (**Figure S2**D).

S4.2.4. ***Rate of phagocytosis of dead cells by macrophages***

The rate macrophages phagocytose dead material is described by

$$\frac{dM_{\Phi E}}{dt} = -d_{D,M\Phi} D M_{\Phi E}, \qquad S31$$

$$\frac{dM_{\Phi F}}{dt} = d_{D,M\Phi} D M_{\Phi E}, \qquad S32$$

$$\frac{dD}{dt} = -d_{D,M\Phi} D (M_{\Phi E} - M_{\Phi F}), \qquad S33$$

where empty macrophages ($M_{\Phi E}$) phagocytose dead cells ($D$) at a rate $d_{D,M\Phi}$ and become loaded macrophages ($M_{\Phi F}$). Loaded macrophages also phagocytose dead cells at a rate $d_{D,M\Phi}$. Assuming the initial concentration of macrophages and dead cells is $M_{\Phi E}(0) = 36 \times 10^7$ cells/ml, $M_{\Phi F}(0) = 0$ cells/ml, and $D(0) = 5 \times 36 \times 10^7$ cells/ml (based on our full model's initial conditions and the experiment where the percentage of macrophages that had engulfed material over 25 hours was measured[27]), we fit the rate macrophages phagocytose dead material over 25 hours and obtained $d_{D,M\Phi} = 8.03$ per cell/day (**Figure S2**E).

S4.2.1. ***Clearance of extracellular virus by macrophages***

To determine the clearance rate of extracellular virus by macrophages, $\delta_{V,M\Phi}$, we used measurements of foot-and-mouth disease virus (FMDV) uptake in macrophages over 2 hours *in vitro*[23] with the simple model

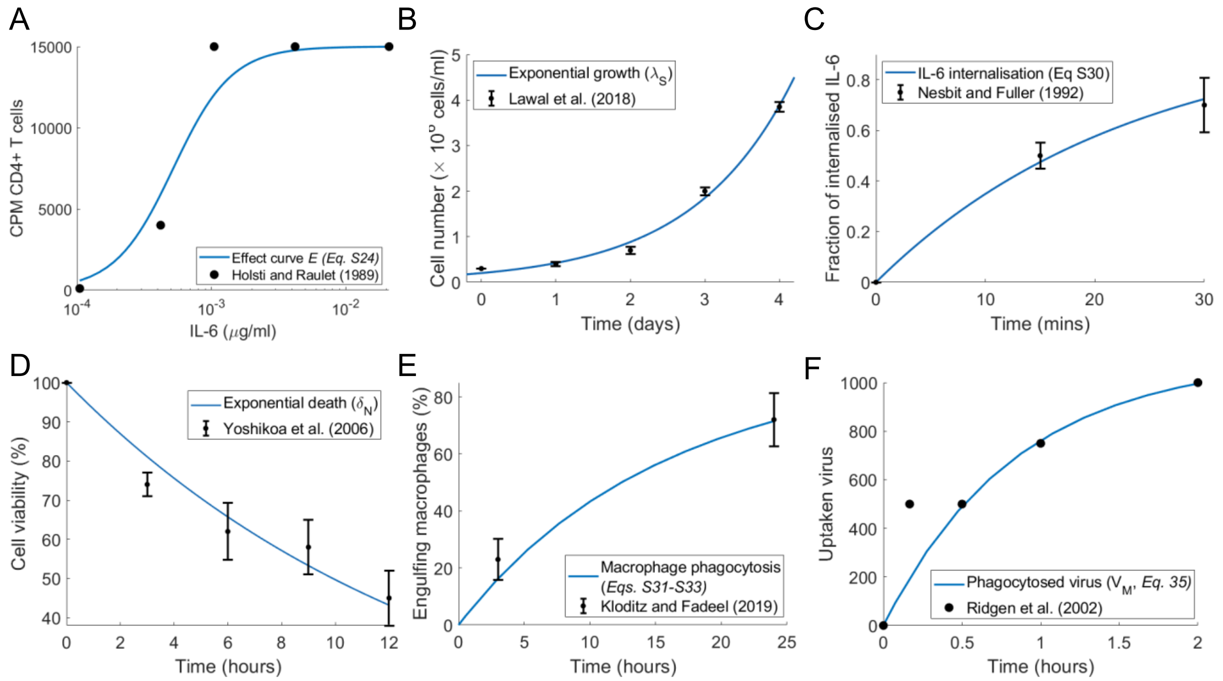


***Figure S2. Dynamics of IL-6 on T cell expansion, epithelial cell growth, IL-6 internalization, neutrophil-induced damage, and macrophage phagocytosis. A)*** *Effect curve* (**Eq. S24**) *for the IL-6 effect on T cell expansion fit to measurements CD4⁺ T cells from dilutions of IL-6 by Holsti and Raulet*[21]. ***B)*** *Exponential growth curve fit to the growth of A549 cells*[2] ***C)*** *The internalization rate of IL-6 (**Eq. S30**) fit to the fraction of internalized IL-6*[47]. ***D)*** *Exponential decay fit to cell viability after $H_2O_2$ administration*[24]. ***E)*** *The macrophage clearance of apoptotic material (**Eqs. S31-S33**) was fit to the percentage of macrophages that had engulfed material over 25 hours*[27]. ***F)*** *The phagocytosis rate of extracellular virus by macrophages was obtained by fitting* **Eqs. S34-S35** *to the uptake of virus by macrophages measured by Rigden et al.*[23]. *Data (black) is plotted as either circles (**A & F**) or mean and standard deviation error bars (**B-E**); solid blue line: corresponding fit.*

$$\frac{dV}{dt} = -\delta_{V,M\Phi} V M_{\Phi I}, \qquad \text{S34}$$

$$\frac{dV_M}{dt} = \delta_{V,M\Phi} V M_{\Phi I}, \qquad \text{S35}$$

where $V$ is free virus, and $V_M$ is the amount of phagocytosed virus. Considering $M_{\Phi I} = 36 \times 10^7$ cells/ml (i.e. our initial measurements in the lung) was constant gave an estimate of $\delta_{V,M\Phi} = 768$/day (**Figure S2**F).

##### S4.2.1. *Production of type I IFN by monocytes*

We modelled the production of type I IFN by monocytes (**Eq. S17**) by

$$\frac{dF_U}{dt} = \frac{p_{F,M} M}{M + \eta_{F,M}}. \qquad \text{S36}$$

To fit the production of IFN from monocytes, we considered a simple production function for monocytes in the absence of any cytokine or inflammatory signalling

$$\frac{dM}{dt} = \frac{p_{\hat{M}} M^h}{M^h + EC_{50,\hat{M}}^h}\left(1 - \frac{M}{\hat{M}_{max}}\right), \qquad \text{S37}$$

where $p_{\hat{M}}$ is the rate of monocyte production per day, $EC_{50,\hat{M}}$ is the production half-effect, $h$ is the Hill coefficient, and $\hat{M}_{max}$ is the carrying capacity of the monocyte population. Ohta et al.[42] measured the number of monocytes after incubation for 12 days with 0.1 nM of calcitriol. Fitting **Eq. S37** to their data, we obtained $EC_{50,\widehat{M}} = 5.4 \times 10^4$ cells, $h = 13.8$, and $\hat{p} = 9.4 \times 10^4$ cells/day (**Figure S3**A).

Krilov et al.[43] measured IFN-α production from monocytes that had been cultured for either 1, 2, 4 or 7 days before the introduction of respiratory syncytial virus (RSV). IFN-α was measured 24 hours after RSV was introduced at $t_{RSV}$. Combining **Eqs. S36-S37**, gives the production of IFN ($F_U$) from RSV stimulation of monocytes

$$\frac{d\widehat{M}}{dt} = \frac{p\widehat{M}^h}{M^h + EC_{50,\widehat{M}}^h}\left(1 - \frac{\widehat{M}}{M_{max}}\right), \quad \text{S38}$$

$$\frac{dF_U}{dt} = \frac{p_{F,M}M^h}{M^h + \eta_{F,M}^h} H(t - t_{RSV}), \quad \text{S39}$$

assuming IFN production occurs only after RVS introduction at $t_{RSV}$=1, 2, 4, or 7 days (modelled using the Heaviside function). Fixing $\eta_{F,M} = EC_{50,\widehat{M}} = 0.54$ pg/ml, and setting the production rate of monocytes to be equivalent to Ohta's experiments (**Figure S3**A), we fit the concentration of IFN 24 hours after $RSV$ is introduced ($F_U(t_{RSV} + 24)$) and obtained $p_{F,M} = 997.1978$ IU/ml/day (**Figure S3**B). Converting the production rate using IFN's specific activity of 0.028 IU/pg gives $p_{F,M} = 3.561$ pg/ml/day

##### S4.2.1. *Resident macrophage production rate during declining infection*

Tissue-resident (or alveolar) macrophages return to homeostasis after viral infections have been successfully cleared[86], which we accounted for using logistic production (**Eqs. S6-S7**):

$$\frac{dV}{dt} = -d_V V, \quad \text{S40}$$

$$\frac{dM_{\Phi R}}{dt} = \frac{\lambda_{M\Phi}M_{\Phi I}}{V + \epsilon_{V,M\Phi}}\left(1 - \frac{M_{\Phi R}}{M_{\Phi max}}\right), \quad \text{S41}$$

$$\frac{dM_{\Phi I}}{dt} = -d_{M\Phi}M_{\Phi I} - \frac{\lambda_{M\Phi}M_{\Phi I}}{V + \epsilon_{V,M\Phi}}\left(1 - \frac{M_{\Phi R}}{M_{\Phi max}}\right). \quad \text{S42}$$

Here $\lambda_{M\Phi}$ and $\epsilon_{V,M\Phi}$ were fit and all other parameters were fixed to their estimated values (**Table S1**).

We infected mice with 75 $TCID_{50}$ influenza A/Puerto Rico/34/8 (PR8) and measured viral loads[87] and alveolar macrophages ($F480^{hi}CD11c^{hi}CD11B^{-}$, see **Figure S3**D-E). Fitting **Eqs. S40-S42** to this data (**Figure S3**D-E) resulted in estimates of $\lambda_{M\Phi} = 0.082$ $TCID_{50}$/day, $\epsilon_{V,M\Phi} = 63.1$ $TCID_{50}$, $M_{\Phi,max} = 5.02$ cells, and $d_V = 1.43$/day. To convert from $TCID_{50}$ to a viral copies (RNA copy number) per volume (ml) for the units in our model, we used correlations between these (**Figure S3**C) for influenza A matrix[88]. We assumed 3.5 $TCID_{50}$ was equivalent to $2.19 \times 10^5$ virus copies[89] and took the ratio between the $TCID_{50}$ /100 µl and $10^6$ copies/100 µl to be approximately 0.37 (**Figure S3**C). Thus, we set $\lambda_{M\Phi} = 5.94 \times 10^3$ copies/ml/day and $\epsilon_{V,M\Phi} = 905.22$ copies/ml in our simulations. We validated these estimates against data from Landsman and Jung[4] (not shown).

##### S4.2.1. *Production of GM-CSF by monocytes*

Lee et al.[40] measured the concentration of GM-CSF produced by adherent monocytes incubated with LPS over 72 hours (**Figure S3**F). To fit to this data, we developed a simplified submodel given by

$$\frac{dM}{dt} = \frac{p_{M,G}G_B^{h_M}}{G_B^{h_M} + \epsilon_{G,M}} - d_M M, \quad \text{S43}$$

$$\frac{dG_U}{dt} = \frac{p_{G,M}M}{M+\eta_{G,M}} - k_{lin_G}G_U - k_{B_G}(MA_G - G_B)G_U + k_{U_G}G_{B,}, \quad S44$$

$$\frac{dG_B}{dt} = -k_{int_G}G_B + k_{B_G}(MA_G - G_B)G_U - k_{U_G}G_B, \quad S45$$

where the production of monocytes ($M$) by bound GM-CSF ($G_B$) was modelled by a Hill function. Parameters were calibrated to homeostasis and we found the production rate of monocytes by GM-CSF to be $p_{M,G} = 7.29 \times 10^3$ cells/ml and the production of GM-CSF by monocytes to be $p_{G,M} = 7.7 \times 10^5$ pg/ml/day through fitting to the GM-CSF measurements of Lee et al.[40] (**Figure S3**F).

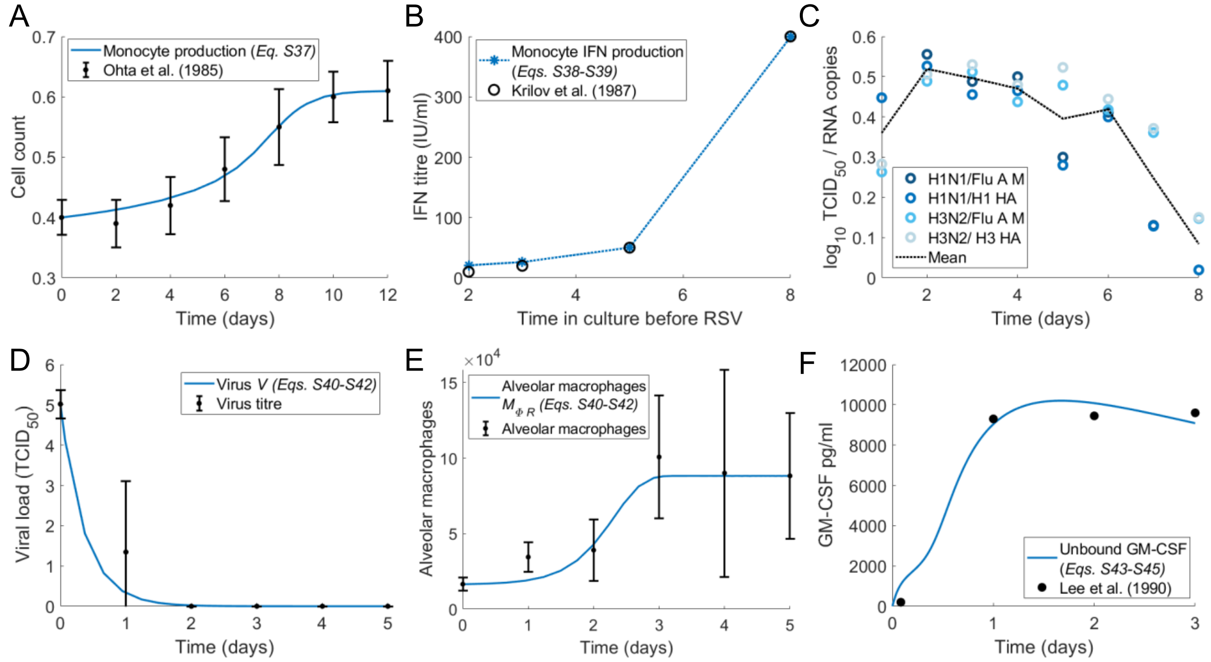


***Figure S3. Monocyte expansion and type I IFN production by monocytes, alveolar macrophage replenishment after viral infection, and GM-CSF production by monocytes. A) Eq. S37** fit to time course of proliferation of monocytes in culture[42]. **B)** Fit of **Eqs. S38-S39** to the production of IFN-α by monocytes after 24 hours with RSV as a function of the number of days of pre-culturing (1, 2, 4 or 7)[43]. **C)** Correlation between infectious virus titre and RT-PCR copy number for influenza A and B measured by Laurie et al.[88] The relative $TCID_{50}$ compared to the RNA copies is plotted for each virus strain and the mean as a black dashed line. **D-E)** Fit of **Eqs. S40-S42** to viral loads[87] and alveolar macrophages from experimental influenza infections. **F)** The production of GM-CSF from stimulated monocytes was recorded by Lee et al.[40] Using a simplified version of the full model (**Eqs. S43-S46**), we obtained the production rates for monocytes and GM-CSF. Data (black) is plotted as either circles/stars (**B&F**) or mean and standard deviation error bars (**A,D-E**); solid blue line: corresponding fit.*

##### S4.2.1. *Production of IFN by infected cells*

To determine the production rate of type I IFN by infected cells, we considered all immune populations to be zero in the full model (**Eqs. S1-S22**), giving

$$\frac{dV}{dt} = pI - d_V V, \quad S46$$

$$\frac{dS}{dt} = \lambda_S\left(1 - \frac{S+I}{S_{max}}\right)S - \beta SV, \quad S47$$

$$\frac{dI}{dt} = \frac{\beta}{1 + F_B/\epsilon_{F,I}} S(t - \tau_I)V(t - \tau_I) - d_I I, \qquad S48$$

$$\frac{dF_U}{dt} = \psi^*_{prod,F} + \frac{p_{F,I} I}{I + \eta_{F,I}} - k_{lin_F} F_U - k_{B_F}(IA_F - F_B)F_U + k_{U_F} F_B, \qquad S49$$

$$\frac{dF_B}{dt} = -k_{int_F} F_B + k_{B_F}(IA_F - F_B)F_U - k_{U_F} F_B. \qquad S50$$

Here IFN is only produced by infected cells and so there is an additional homeostatic production of IFN, $\psi^*_{prod,F}$, to account for general macrophage and monocyte production. $\psi^*_{prod,F}$ is obtained from calculating homeostasis for $F_U$ and $F_B$, i.e. $dF_U/dt = dF_B/dt = 0$. Resistant cells ($R$) were not considered in this model as the data was only measured over 1 day. By fixing all parameters to their previously established values (**Table S1**), and fitting $p_{F,I}$ and $\eta_{F,I}$, we obtained $p_{F,I} = 2.823 \times 10^4$ pg/ml/day and $\eta_{F,I} = 0.00112$ pg/ml (**Figure S4**A). Since the concentration of IFN-$\alpha$ in patients infected with SARS-CoV-2 is lower than IFN-$\beta$, we reduced the production rate to $p_{F,I} = 2.823$ pg/ml/day so that model dynamics lay within the ranges of IFN-$\alpha$ exhibited by patients with SARS-CoV-2 infection (**Figure S6**A-B and Laing et al.[80]).

##### S4.2.2. *Production of IL-6 by infected cells*

To determine the rate of production of IL-6 by infected cells, Ye *et al*[36] measured the *in vitro* replication kinetics of H5N1 and H7N9 viruses in A549 cells. Cells were infected by either virus at an MOI of 2 and grown to confluence in sterile T75-tissue culture flasks (approximate cell number of $8.4 \times 10^6$)[90] and the concentration of IL-6 released from A549 cells in response to infection with both viruses was measured. We reduced the full model (**Eqs. S1-S22**) to only consider virus infection and IL-6 production by infected cells

$$\frac{dV}{dt} = pI - d_V V, \qquad S51$$

$$\frac{dS}{dt} = -\beta SV, \qquad S52$$

$$\frac{dI}{dt} = \beta SV - d_I I, \qquad S53$$

$$\frac{dL_U}{dt} = \frac{p_{L,I} I}{I + \eta_{L,I}}. \qquad S54$$

and fit to this data to obtain $p_{L,I} = 11.887$ pg/ml and $\eta_{L,I} = 0.7232 \times 10^9$ cells/ml (**Figure S4**B-C).

##### S4.2.3. *Production of IL-6 by alveolar macrophages*

We modelled the production of IL-6 by alveolar macrophages by

$$\frac{dL_U}{dt} = \frac{p_{L,M_{\Phi I}} M_{\Phi \mathrm{I}}}{M_{\Phi I} + \eta_{L,M\Phi}} \qquad S55$$

to compare to observations from Shibata et al.[37] who measured the production of IL-6 by alveolar macrophages stimulated by different concentrations of LPS. Assuming no proliferation of macrophages from LPS introduction but that LPS scales the production of IL-6, we modified the above equation to be

$$\frac{dL_U}{dt} = \frac{p_{L,M_{\Phi I}} M_{\Phi \mathrm{I}}}{M_{\Phi I} + \eta_{L,M\Phi}} LPS, \qquad S56$$

and fit the Shibata et al. data and obtained $p_{L,M_{\Phi I}} = 0.078$ ng/ml/hour and $\eta_{L,M\Phi} = 4.47 \times 10^5$ cells/ml (**Figure S4**D).

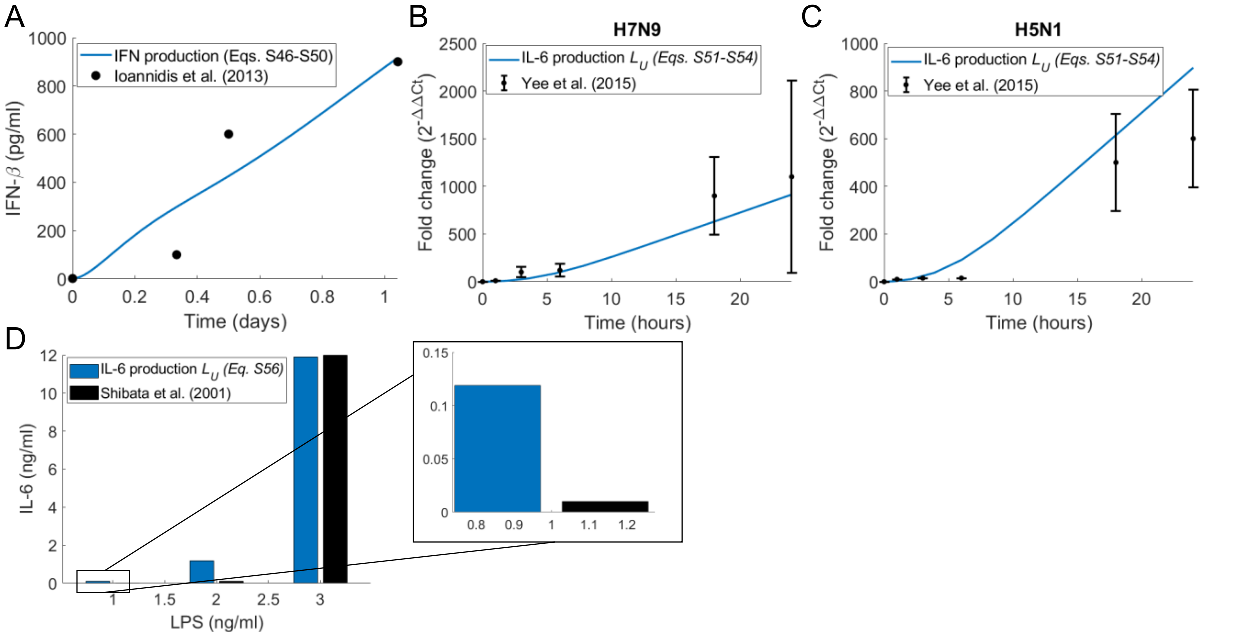


***Figure S4. Production of IFN and IL-6 by infected cells and macrophages. A)*** *Concentration of IFN-β released by alveolar epithelial cells in response to stimulation with influenza virus recorded at 8, 16 and 24 hours*[41]. ***B-C)*** *IL-6 production by infected cells in response to* ***A)*** *H5NA and* ***B)*** *H7N9, measured by Ye et al.*[36] *Data (black) is plotted as mean and standard deviation error bars with the corresponding fit (***Eqs. S51-S54***) in solid blue.* ***D)*** *IL-6 production by macrophages (***Eq. S56***) in response to stimulation with LPS of varying dosage sizes. Shibata et al.* [37] *measured the production of IL-6 for different dosages of LPS and fitting the production rate to this data to obtain* $p_{L,M\Phi}, \eta_{L,M\Phi}$.

#### S5. Parameters calculated from homeostasis

Remaining parameters in the model were estimated to ensure the model maintained homeostasis in the absence of infection, i.e. we required the system to return to equilibrium state after small perturbations in initial conditions for the immune cells and cytokines. Homeostasis equations are defined below (**Eqs. S57-S70**), along with the corresponding parameter they define. These were determined by solving $d/dt = 0$. At homeostasis we assume there to be no virus and resistant cells ($V = R = 0$). Here $X^*$ represents homeostatic values.

$$F_B(0) = F_B^* = \frac{k_{B_F} T^* A_F F_U^*}{k_{int_F} + k_{B_F} F_U^* + k_{U_F}}, \qquad S57$$

$$G_B(0) = G_B^* = \frac{k_{B_G} M^* A_G G_U^*}{k_{int_G} + k_{B_G} G_U^* + k_{U_G}}, \qquad S58$$

$$C_B(0) = C_B^* = \frac{k_{B_C} {C_U^*}^{POW_C} A_C N^*}{k_{int_C} + k_{B_C} {C_U^*}^{POW_C} + k_{U_C}}, \qquad S59$$

$$C_{BF}(0) = C_{BF}^* = \frac{C_B^*}{A_C N^*}, \qquad S60$$

$$L_B(0) = L_B^* = \frac{k_{B_L}(T^* + N^* + M^*) A_L L_U^*}{k_{int_L} + k_{B_L} L_U^* + k_{U_L}}, \qquad S61$$

$$M_{\Phi I}(0) = M_{\Phi I}^* = \frac{\left(\frac{p_{M_{\Phi \mathrm{I}},G} {G_B^*}^{h_{M,M\Phi}} M^*}{{G_B^*}^{h_{M,M\Phi}} + {\epsilon_{G,M\Phi}}^{h_{M,M\Phi}}} + \frac{p_{M_{\Phi I},L} L_B^* M^*}{L_B^* + \epsilon_{L,M\Phi}}\right)}{\left(1 - \frac{M_{\Phi R}^*}{M_{\Phi max}}\right)\frac{\lambda_{M\Phi}}{\epsilon_{V,M\Phi}} + d_{M_{\Phi \mathrm{I}}}}, \quad S62$$

$$\eta_{C,M} = \frac{p_{C,M} M^* - M^*\left(k_{lin_C} C_U^* + k_{B_C}(N^* A_C - C_B^*){C_U^*}^{POW_C} - k_{U_C} C_B^*\right)}{k_{lin_C} C_U^* + k_{B_C}(N^* A_C - C_B^*){C_U^*}^{POW_C} - k_{U_C} C_B^*}, \quad S63$$

$$p_{L,M\Phi} = \frac{M_{\Phi I}^* + \eta_{L,M\Phi}}{M_{\Phi I}^*}\left(-\frac{p_{L,M} M^*}{M^* + \eta_{L,M}} + k_{lin_L} L_U^* + k_{B_L}\left((N^* + T^* + M^*)A_L - L_B^*\right)L_U^* - k_{U_L} L_B^*\right), \quad S64$$

$$p_{G,M\Phi_I} = \frac{k_{lin_G} G_U^* + k_{B_G}(M^* A_G - G_B^*)G_U^* - k_{U_G G_B^*}}{\frac{M_{\Phi I}^*}{M_{\Phi I}^* + \eta_{G,M\Phi}} + \frac{M^*}{M^* + \eta_{G,M}}}, \quad S65$$

$$p_{M,G} = \left(\frac{{G_B^*}^{h_M} + {\epsilon_{G,M}}^{h_M}}{{G_B^*}^{h_M}}\right)\left(\frac{p_{M_{\Phi \mathrm{I}},G} {G_B^*}^{h_{M,M\Phi}} M^*}{{G_B^*}^{h_{M,M\Phi}} + {\epsilon_{G,M\Phi}}^{h_{M,M\Phi}}} + \frac{p_{M_{\Phi I},L} L_B^* M^*}{L_B^* + \epsilon_{L,M\Phi}} + d_M M^*\right), \quad S66$$

$$\eta_{F,M\Phi} = \frac{p_{F,M\Phi} M_{\Phi \mathrm{I}}^* + \left(\frac{p_{F,M} M^*}{M^* + \eta_{F,M}} - k_{lin_F} F_U^* - k_{B_F}(T^* A_F - F_B^*)F_U^* + k_{U_F} F_B^*\right) M_{\Phi I}^*}{-\frac{p_{F,M} M^*}{M^* + \eta_{F,M}} + k_{lin_F} F_U^* + k_{B_F}(T^* A_F - F_B^*)F_U^* - k_{U_F} F_B^*}, \quad S67$$

$$T_{prod}^* = d_T T^* - \frac{p_{T,L} L_B^* T^*}{L_B^* + \epsilon_{L,T}} - \frac{p_{T,F} F_B^* T^*}{F_B^* + \epsilon_{F,T}}, \quad S68$$

$$N_{prod}^* = \left(d_N N^* - \frac{p_{N_L} L_B^*}{L_B^* + \epsilon_{L,N}}\right)\frac{1}{NR}, \quad S69$$

$$M_{prod}^* = \frac{\frac{1}{MR}\left(\frac{p_{M_{\Phi I},G} {G_B^*}^{h_{M,M\Phi}} M^*}{{G_B^*}^{h_{M,M\Phi}} + {\epsilon_{G,M\Phi}}^{h_{M,M\Phi}}} + \frac{p_{M_{\Phi I},L} L_B^* M^*}{L_B^* + \epsilon_{L,M}} + d_M M^*\right) - \psi_M^{max}\frac{{G_B^*}^{h_M}}{{G_B^*}^{h_M} + {\epsilon_{G,M}}^{h_M}}}{1 - \frac{{G_B^*}^{h_M}}{{G_B^*}^{h_M} + {\epsilon_{G,M}}^{h_M}}}, \quad S70$$

The model was then simulated to confirm the parameter values determined resulted in a stable system at homeostasis (**Figure S5**).

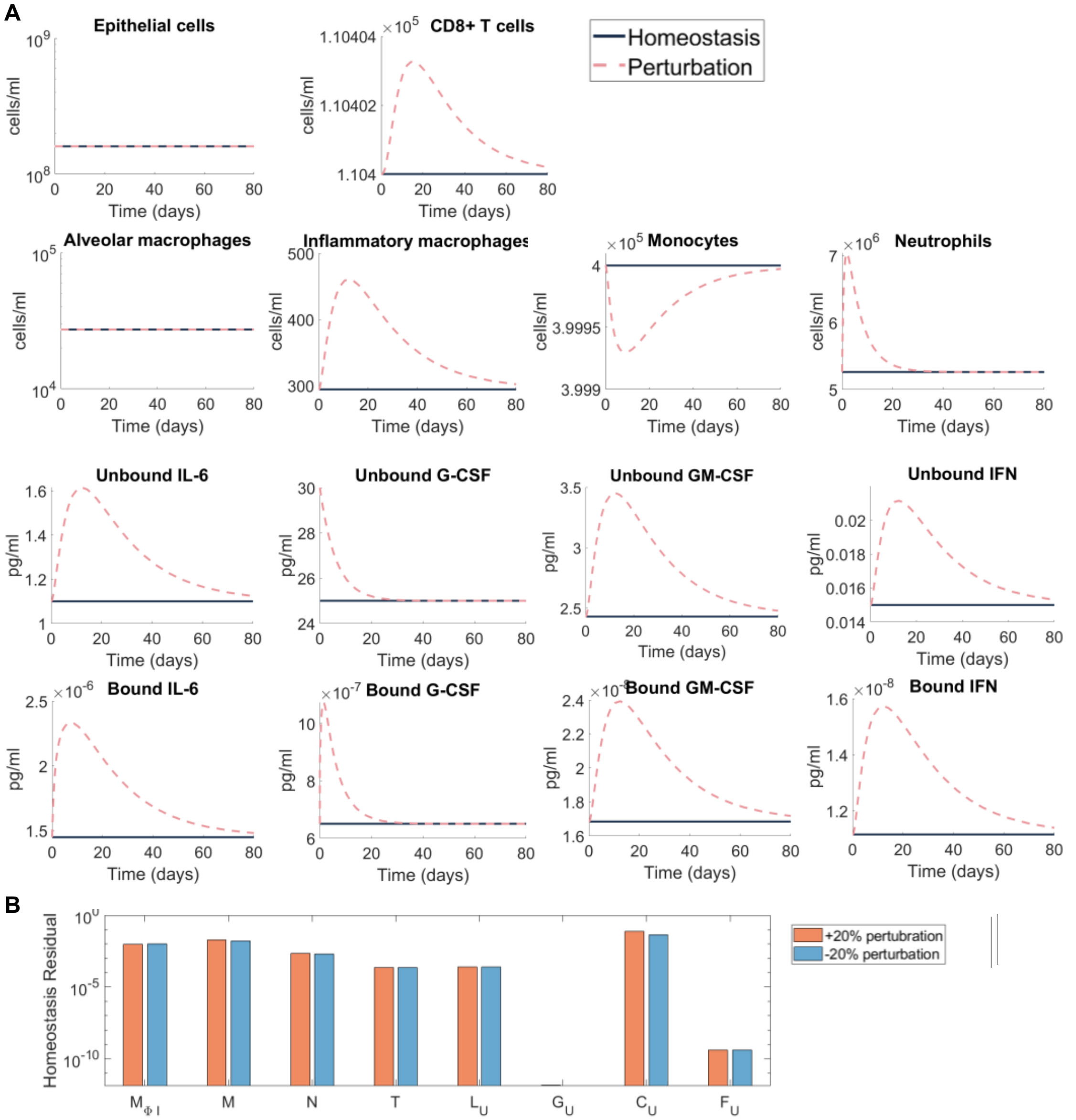


***Figure S5. Homeostatic disease-free system regulation. A)*** *To confirm that parameters in the model represented realistic immunocompetent individuals in the disease-free scenario,* ***Eqs. S1-S22*** *were simulated where* $V_0 = 0$ *and parameters were given by the homeostasis* ***Eqs. S57-S70****. The initial concentration of G-CSF was perturbed and compared to simulations of the model at homeostasis. Simulations at homeostasis are represented by solid lines (purple) and perturbed simulations as dashed lines (pink).* ***B)*** *The maximum residual between variables and their initial conditions at day 50 was measured to confirm that the system was stable for perturbations in all immune cells and cytokines.*

#### S6. Model predication validation and sensitivity analysis

Having confirmed that predicted dynamics from the reduced IFN model (**Eqs. 31-37** and **Figure. 3**, **Main Text**) qualitatively matched to time-series measurements for IFN-$\alpha$ from Trouillet-Assant et al.[70] (**Figure S6**A), we next sought to further validate predictions of the full model by comparing IFN, IL-6, and G-CSF dynamics to previously published observations (**Figure S6**B-F). For this, measurements for IFN-$\alpha 2$ and IL-6 plasma concentrations were taken from Trouillet-Assant et al.[70] (**Figure S6**B-C), and IL-6 concentrations from patients requiring and not requiring mechanical ventilation obtained by Herold et al.[91] were also used to validate predicted IL-6 dynamics (**Figure S6**D). IL-6 and G-CSF plasma concentrations in symptomatic and asymptomatic patients were obtained by Long et al.[92] (**Figure S6**E-F). Full model simulations are given in **Figure S7**.

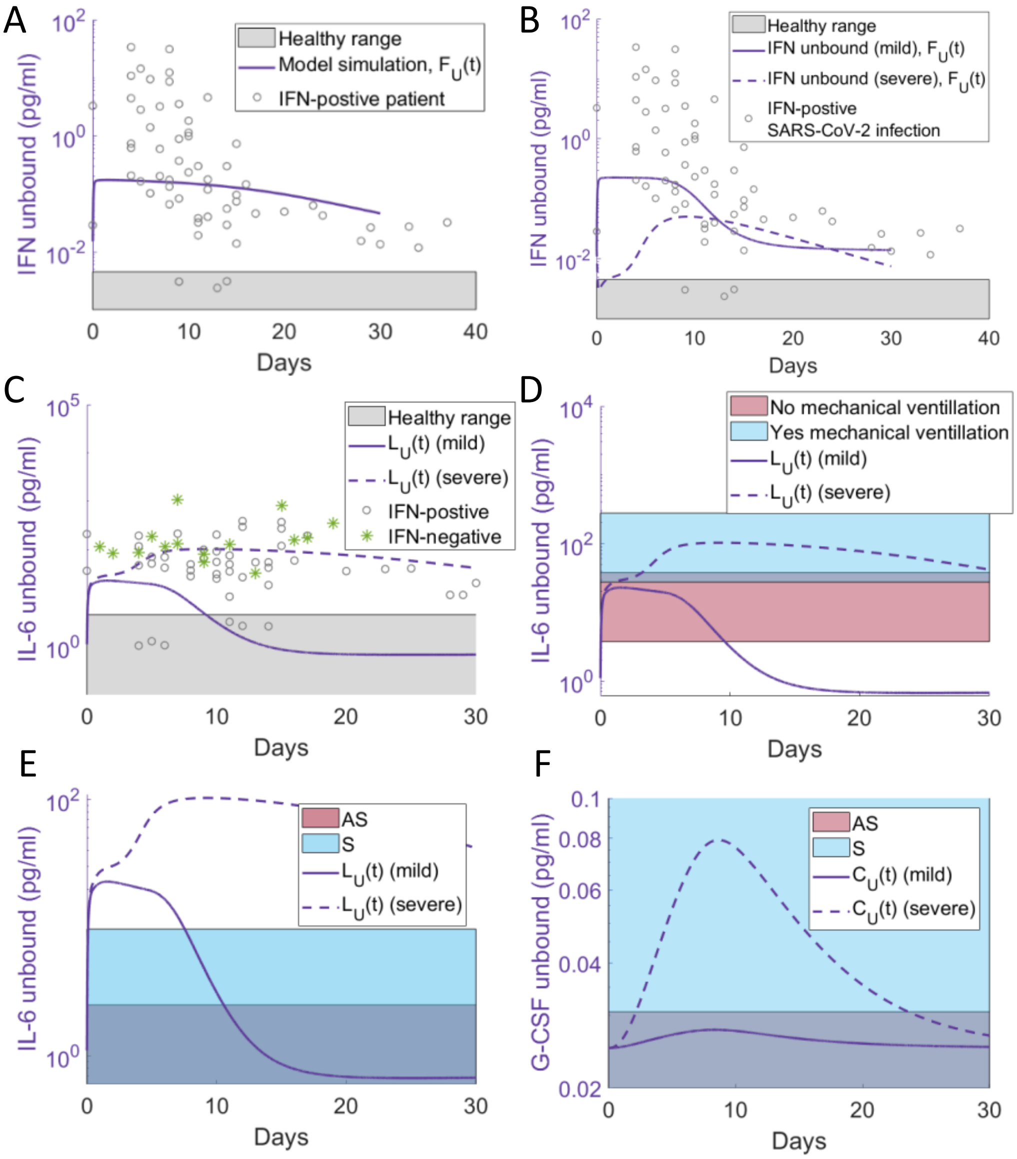


***Figure S6. Model validation against human cytokine measurements during SARS-CoV-2 infection. A)*** *IFN dynamics of the reduced model (**Figure 3 Main Tex**t) overlaid with patient IFN-$\alpha$2 plasma concentrations from Trouillet-Assant et al.*[70] *The solid line (purple) represents the unbound IFN dynamics from the reduced model **(Eqs. 27-33).** Individual patient IFN-$\alpha$2 measurements are plotted as grey circles. Normal IFN-$\alpha$2 concentration in healthy volunteers are indicated by a grey area. **B-F)** Mild and severe dynamics (**Eqs. S1-S22**) corresponding to simulations in **Figure 4 Main Text** and **Figure S7** overlaid with measurements from the literature with solid lines: mild disease dynamics; dashed lines: severe disease dynamics.. **B-C)** Plasma IFN-$\alpha$ and IL-6 in COVID-19 critically ill patients (n=26) obtained by Trouillet-Assant et al.*[70] *overlaid with mild and severe unbound IFN ($F_U(t)$) and mild and severe unbound IL-6 ($L_U(t)$). **D)** IL-6 levels in patients requiring and not requiring mechanical ventilation obtained by Herold et al.*[91] *overlaid with mild and severe unbound IL-6 dynamics. **E-F)** IL-6 and G-CSF plasma concentration obtained by Long et al.*[92] *in symptomatic "S" and asymptomatic "AS" COVID-19 patients overlaid with corresponding mild and severe model dynamics.*

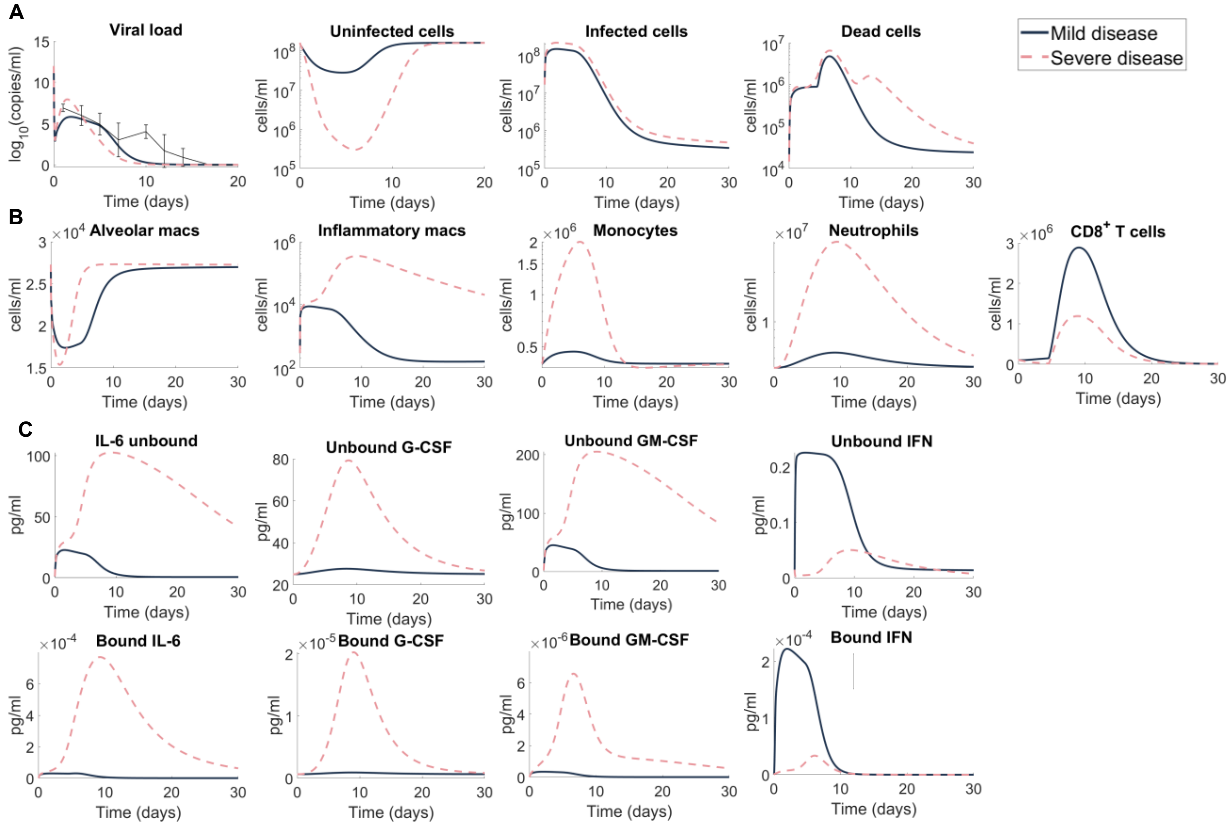


***Figure S7. Predicting mild and severe COVID-19 dynamics (all model variables).*** *Extension of results of mild and severe disease dynamics in* ***Figure 4 Main Text****. Mild disease (solid lines) dynamics obtained by using baseline parameter estimates (****Tables S1****) while severe disease dynamics (dashed lines) were obtained by decreasing the production rate of type I IFN,* $p_{F,I}$*, and increasing the production of monocytes,* $p_{M,I}$*, and their differentiation to macrophages,* $\eta_{F,M\Phi}$*.* ***A)*** *Lung cells concentrations (susceptible cells* $S(t)$*, resistant cells* $R(t)$*, infected cells* $I(t)$*, dead cells* $D(t)$ *and virus* $V(t)$*). Solid black line with error bars indicates macaque data (see* ***Fig. 2 Main Text****).* ***B)*** *Immune cell concentrations (resident macrophages* $M_{\Phi R}(t)$*, inflammatory macrophages* $M_{\Phi I}(t)$*, monocytes* $M(t)$*, neutrophils* $N(t)$ *and T cells* $T(t)$*).* ***C)*** *Bound and unbound cytokine concentrations (IL-6 unbound* $L_U(t)$ *and bound* $L_B(t)$*, GM-CSF unbound* $G_U(t)$ *and bound* $G_B(t)$*, G-CSF unbound* $C_U(t)$ *and bound* $C_B(t)$*, type I IFN unbound* $F_U(t)$ *and bound* $F_B(t)$*).*

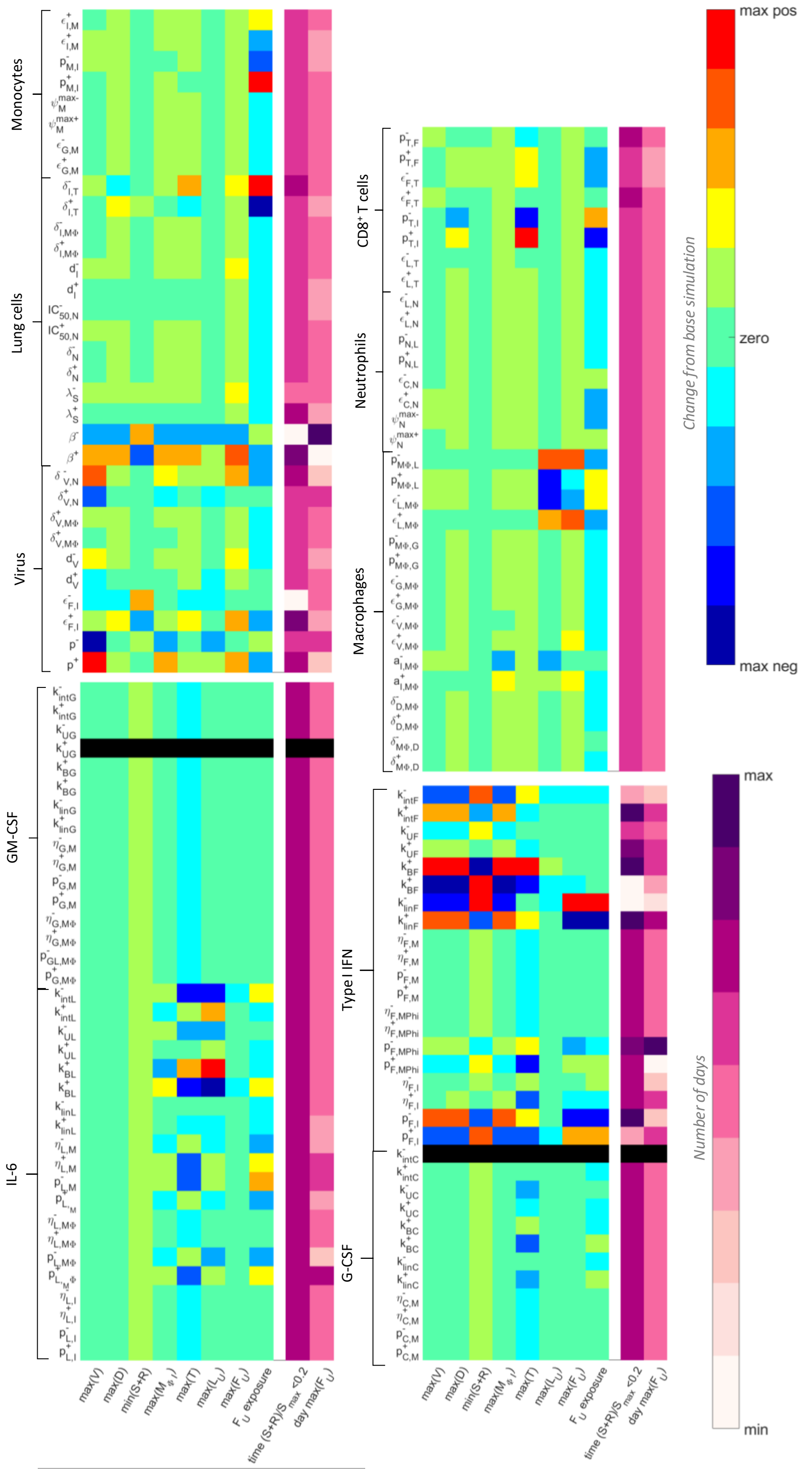


***Figure S8. Full analysis of parameters driving COVID-19 severity.*** *A local sensitivity analysis was performed by varying each parameter ±20% from its originally estimated value and simulating the model. Predictions were then compared to baseline considering: Maximum viral load (max(V)), maximum concentration of dead cells (max(D)), minimum uninfected live cells (min(S+R)), maximum concentration of inflammatory macrophages (max($M_{\Phi I}$)), maximum number of* CD8$^{+}$ *T cells (max(T)), maximum concentration of IL-6 (max($L_U$)), maximum concentration of type I IFN (max($F_U$)), the total exposure to type I IFN ($F_U$ exposure), the number of days damaged tissue was >80% (time $(S+R)/S_{max}$)<0.2), and the day type I IFN reached its maximum (day max($F_U$)). The heatmaps show the fold change of each metric, where blue signifies the minimum value observed and red signifies the maximum value observed, or by the number of days, where light to dark pink signifying increasing number of days from zero. The most sensitive parameters are shown in* ***Figure 5*** *in the* ***Main Text.***

#### *S6.1. Generating of virtual patients*

Initial parameter sets for each virtual patient were drawn from normal distributions with means fixed to the corresponding parameter value in **Table S1** and standard deviations derived from appropriate standard deviation or confidence interval measurements in the literature. Specifically, the standard deviation for

- the half-effect concentration of IFN on viral infectivity ($\epsilon_{F,I}$) was informed by the 95% confidence interval from fitting the Emax curve to MERS-CoV-expression nanoluciferase (nLUC) reported by Sheahan et al.[15], from the IFN-$\alpha$ 95% confidence interval on day 0 from Trouillet-Assant et al.[70] (**Figure S6**B-C),
- IFN production by infected cells ($p_{F,I}$ and $\eta_{F,I}$) and IFN production by macrophages ($\eta_{F,M\Phi}$) were drawn from IL-6 concentration in no mechanical ventilation patients (mild) and mechanical ventilation patients (severe) from Herold et al.[91],
- the production of IL-6 by macrophages and macrophages by IL-6 ($p_{L,M\Phi}$, and $p_{M\Phi,L}$) from Liu et al. [93] (**Figure S6**D), and
- the production of monocytes by infected cells ($p_{M,I}$), and from 95% confidence interval generated from estimating the parameter for production of IFN by monocytes[42] ($p_{F,M}$; **Figure S3**).

From normal distributions with standard deviation described above and mean as the original parameter values ($\hat{p}$), we then generated normal distributions covering 99.7% of values lying with 3 standard deviations of the mean, i.e., $[\mu - 3\sigma, \mu + 3\sigma]$[94].

After drawing an initial patient parameter set for each patient, we next used simulated annealing to determine a parameter set that resulted in patient dynamics within physiological ranges[95] for $[l_i, u_i]$ by minimising **Eq. 15**, where $l_i$ and $u_i$ are the upper and lower bounds extracted from measurements for viral load, type I IFN, G-CSF, and IL-6 (**Figure 7 Main Text**). The resulting parameter set from this optimization was then considered to represent a realistic patient and they were accepted into the cohort. The viral, IFN, IL-6, and G-CSF dynamics of the cohort are seen in **Figure S9**, with the physiological ranges used for optimization.

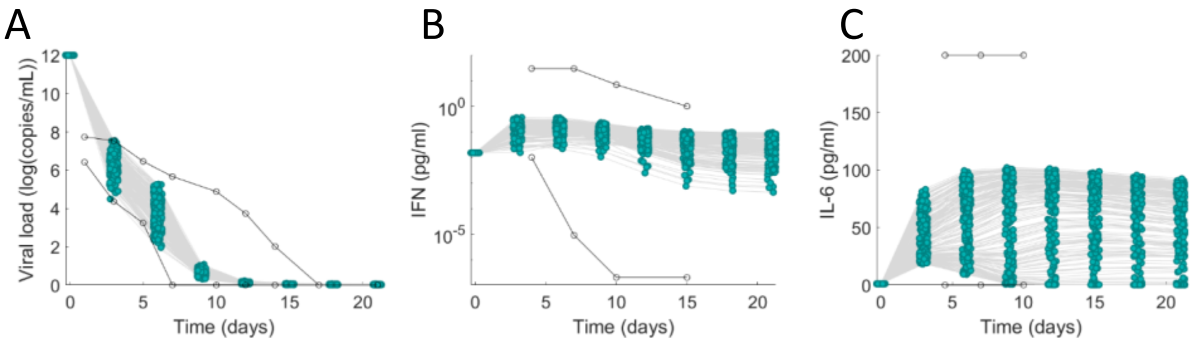


***Figure S9. Cohort dynamics within physiological ranges.*** *Virtual patients were generated so that viral load, IFN and IL-6 concentration were within physiological ranges obtained in the literature. The physiological ranges (denoted by open circles) were obtained from **A)** Munster et al.[96], **B)** Trouillet-Assant et al. [70], and **C)** Herold et al. [91]. Patient dynamics at discrete time points are plotted as joined green dots.*
